# The NS1 protein of influenza B virus binds 5’-triphosphorylated dsRNA to suppress RIG-I activation and the host antiviral response

**DOI:** 10.1101/2023.09.25.559316

**Authors:** Ryan Woltz, Brandon Schweibenz, Susan E. Tsutakawa, Chen Zhao, LiChung Ma, Ben Shurina, Gregory L. Hura, Rachael John, Sergey Vorobiev, GVT Swapna, Mihai Solotchi, John A. Tainer, Robert M. Krug, Smita S. Patel, Gaetano T. Montelione

**Author notes:** Corresponding authors: Gaetano T. Montelione, Center for Biotechnology and Interdisciplinary Sciences, Rensselaer Polytechnic Institute, 1623 15th St, Troy, NY 12180, USA and Smita S. Patel, Department of Biochemistry and Molecular Biology, Robert Wood Johnson Medical School, Rutgers, The State University of New Jersey, Piscataway, NJ, 08854, USA. These two investigators made equal contributions to this project (Ryan Woltz, Brandon Schweibenz).

## Abstract

Influenza A and B viruses overcome the host antiviral response to cause a contagious and often severe human respiratory disease. Here, integrative structural biology and biochemistry studies on non-structural protein 1 of influenza B virus (NS1B) reveal a previously unrecognized viral mechanism for innate immune evasion. Conserved basic groups of its C-terminal domain (NS1B-CTD) bind 5’- triphosphorylated double-stranded RNA (5’ppp-dsRNA), the primary pathogen-associated feature that activates the host retinoic acid-inducible gene I protein (RIG-I) to initiate interferon synthesis and the cellular antiviral response. Like RIG-I, NS1B-CTD preferentially binds blunt-end 5’ppp-dsRNA. NS1B-CTD also competes with RIG-I for binding 5’ppp-dsRNA, and thus suppresses activation of RIG-I’s ATPase activity. Although the NS1B N-terminal domain also binds dsRNA, it utilizes a different binding mode and lacks 5’ppp-dsRNA end preferences. In cells infected with wild-type influenza B virus, RIG-I activation is inhibited. In contrast, RIG-I activation and the resulting phosphorylation of transcription factor IRF-3 are not inhibited in cells infected with a mutant virus encoding NS1B with a R208A substitution it its CTD that eliminates its 5’ppp-dsRNA binding activity. These results reveal a novel mechanism in which NS1B binds 5’ppp-dsRNA to inhibit the RIG-I antiviral response during influenza B virus infection, and open the door to new avenues for antiviral drug discovery.

## INTRODUCTION

Influenza A and B viruses cause a contagious respiratory disease in humans, resulting in 290,000 to 650,000 deaths worldwide each year (World Health Organization (2008) http://www.who.int/ ), making detailed understanding of their ability to disrupt innate immunity of great biological and biomedical importance. Influenza B virus infections contribute to this death toll, particularly by causing children deaths. The genomes of these negative strand RNA viruses are comprised of eight segments (Krug and Fodor, 2013). The 3’ and 5’ ends of each RNA segment are complementary, forming a dsRNA “panhandle” which when packaged into the virion contain 5’ triphosphorylated (5’ppp) blunt ends (Desselberger et al., 1980; Hsu et al., 1987; Cheong et al., 1999; Bae et al., 2001).

These 5’ppp blunt-ended dsRNA ‘panhandles’ serve as pathogen-associated molecular patterns (PAMPs) for the retinoic acid-inducible gene I (RIG-I/DDX58) protein, a cytosolic pattern recognition receptor responsible for the type-1 interferon response (Sumpter et al., 2005). Synthetic influenza genomic panhandle 5’ppp-dsRNA has been shown to activate RIG-I (Liu et al., 2015). Binding of 5’ppp blunt-ended dsRNA by RIG-I causes structural rearrangements and activates its ATPase and helicase activities (Civril et al., 2011; Jiang et al., 2011; Kowalinski et al., 2011; Luo et al., 2011; Devarkar et al., 2016; Devarkar et al., 2018), initiating an innate immune response that includes phosphorylation and activation of the interferon regulatory factor 3 (IRF3) transcription factor, and subsequent expression of type-I interferons alpha and beta, along with interferon-stimulated genes. This pathway is an important innate immune response to viral infection (Yoneyama et al., 2004; Sumpter *et al*., 2005; Schlee et al., 2009; Poeck et al., 2010; Liu *et al*., 2015).

The smallest genome segments of both influenza A virus A (IAV) and influenza B virus (IBV) viruses encode the non-structural protein NS1, which is crucial in combating host antiviral activities (Krug and Garcia-Sastre, 2013). NS1 is comprised of two structural domains, an N-terminal domain (NTD) and a C-terminal domain (CTD), connected by a flexible polypeptide linker (Krug and Garcia-Sastre, 2013). Most previous studies have focused on the NS1 protein of influenza A virus (NS1A protein), and consequently many functions of the NS1A protein in countering host antiviral functions have been identified and characterized (Krug and Garcia-Sastre, 2013). In particular, dsRNA-binding functions of the N-terminal domains (NTDs) of NS1s from IAV and IBV have been long recognized (Qian et al., 1995; Wang et al., 1999; Donelan et al., 2004; Yin et al., 2007).

More recently, we described the three-dimensional structure and unanticipated RNA-binding properties of the CTD of the NS1 protein from IBV (NS1B-CTD) (Ma et al., 2016). The structure of NS1B-CTD revealed a surface of highly-conserved basic residues, suggesting an RNA-binding function (Ma *et al*., 2016). We showed that NS1B-CTD binds RNA using this strongly conserved surface (Ma *et al*., 2016). Despite the high structural similarity of NS1 CTDs of influenza A and B, this second RNA binding site of IBV NS1s is not shared by NS1s of influenza A viruses, and the CTD of NS1A does not bind RNA substrates (Ma *et al*., 2016). Influenza B viruses containing structure-preserving amino-acid substitutions in this CTD RNA-binding site are attenuated, demonstrating that this RNA-binding activity of NS1B is crucial for influenza B virus replication (Ma et al., 2016). However, the specific role of this RNA-binding during virus infection has not been elucidated.

Here we apply integrative structural biology methods to determine the 3D structures of the complexes formed between the NS1B-CTD and dsRNA molecules. Importantly, these structures provide key clues to the function of this interaction in IBV infection. In these complexes the NS1B-CTD was found to bind to the blunt end of dsRNA. RNA binding studies show that NS1B-CTD preferentially binds 5’ppp dsRNA, 440-fold more tightly than the corresponding non-phosphorylated hairpin dsRNA. Combined circular dichroism (CD), NMR, and small angle X-ray scattering (SAXS) data further demonstrate that NS1B-CTD binds to the blunt-end of 5’ppp-dsRNA, with interactions between the 5’ terminal phosphate and basic residues of NS1s that are strongly conserved across IBV strains, but not in NS1As from IAV strains. The blunt-end binding mode of dsRNA in these NS1B complexes is analogous to that observed in the complex formed between synthetic 5’ppp-dsRNAs and RIG-I (Lu et al., 2010; Jiang *et al*., 2011; Luo et al., 2012; Devarkar *et al*., 2016). Both NS1B-CTD and full-length NS1B compete with 5’ppp dsRNA *in vitro* to suppress RIG-I activation. To establish that NS1B also inhibits RIG-I activation in IBV-infected cells, we also measured phosphorylation of the IRF-3 transcription factor which occurs only when RIG-I is activated. We show that IRF-3 phosphorylation is inhibited in cells infected with the wild-type virus, but is not inhibited in cells infected with a mutant virus encoding a NS1B protein with an amino acid substitution in its CTD that eliminates 5’ppp-dsRNA binding. We conclude that the CTD of NS1B binds 5’ppp-dsRNA to inhibit RIG-I activation during influenza B virus infection.

## RESULTS

### NS1B-CTD binds 5’OH-dsRNA at its blunt ends

The X-ray crystal structure of C-terminal domain of NS1B, amino-acid residues 141-281 (NS1B-CTD), was previously determined at 2.0-Å resolution (Ma *et al*., 2016). NS1B-CTD binds a 5’OH dsRNA substrate used for previous influenza NS1 structural studies [*viz*, (CCAUCCUCUACAGGCG (sense) and CGCCUGUAGAGGAUGG (antisense)] with a K_d_ of ∼ 130 ± 30 nM (Ma *et al*., 2016). Sequence-specific NMR assignments of NS1B-CTD, determined using standard triple-resonance NMR methods, were used to map chemical shift perturbations (CSPs) due to dsRNA binding (Ma *et al*., 2016) onto the NS1B-CTD structure (**Figure 1A)**. These CSPs are all localized to one face of NS1B-CTD. This face includes several surface-exposed basic residues shown to be important for dsRNA binding activity (Ma *et al*., 2016). Little or no chemical shift changes are observed on the opposite face of the molecule, indicating that the overall structure of NS1B-CTD is not changed significantly upon complex formation with dsRNA. Efforts to crystallize this complex were unsuccessful, but yielded an X-ray crystal structure of the dsRNA substrate at 2.3 Å resolution (**Supplementary Figure S1; Supplementary Table S1**). This duplex RNA molecule forms a right-handed, *A*-form, double-stranded helix with 16 base-pairs, and well-defined blunt ends. All nucleotides, including all backbone atoms, were clearly visible in the electron density.

**Fig. 1.**
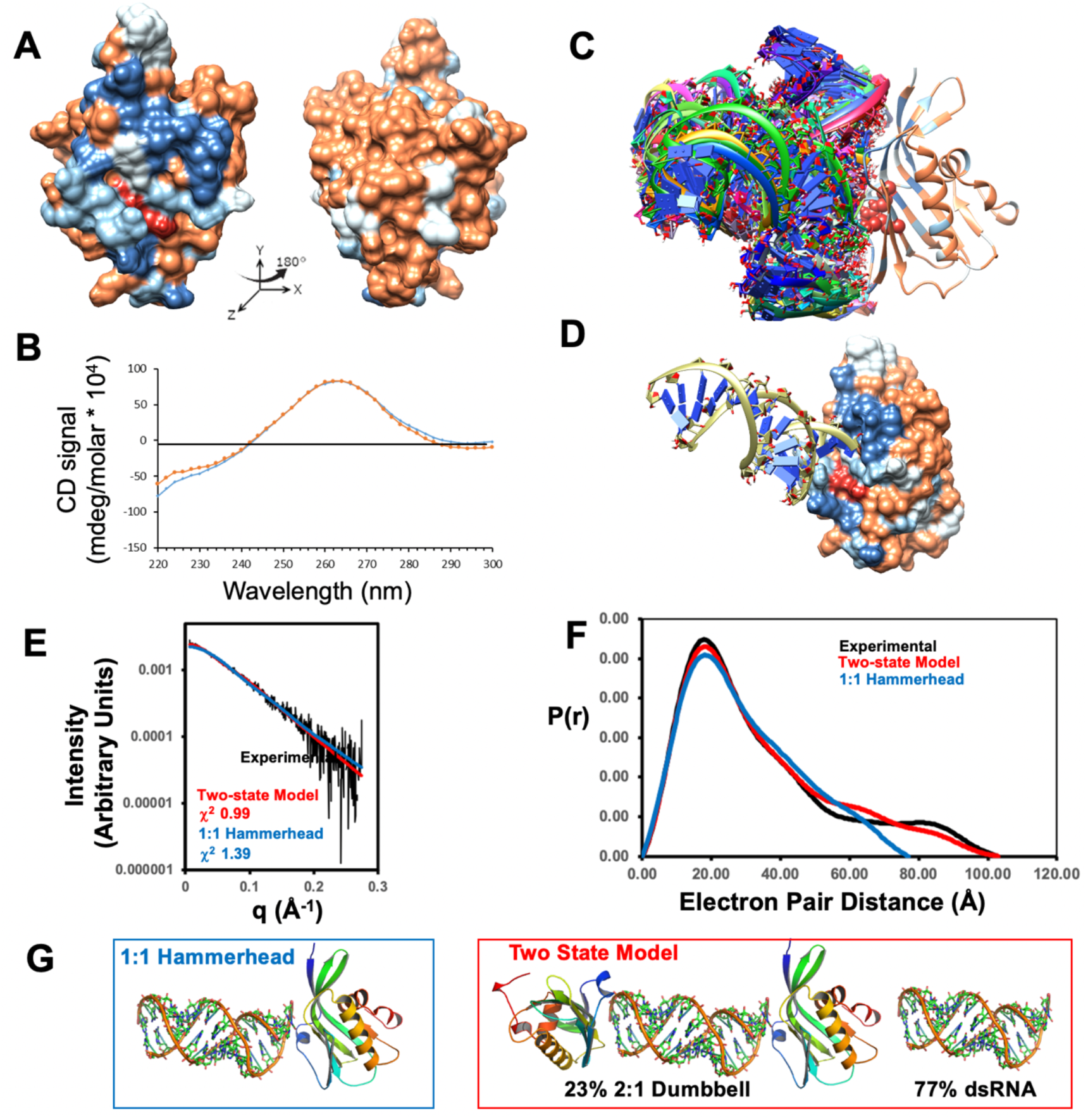
NS1B-CTD* binds 16-bp 5’OH-dsRNA in a blunt end orientation. (A) ^15^N-^1^H chemical shift perturbation (CSP) data due to 16-bp dsRNA binding mapped onto the 3D structure NS1B-CTD (Ma *et al*.). Surface residues are colored to indicate CSPs Δδ_ΝΗ_ > 30 ppb (dark blue), 30 > Δδ_ΝΗ_ > 20 ppb (light blue), or Δδ_ΝΗ_ < 20 ppb (brown). Residues for which Δδ_ΝΗ_ values could not be determined (e.g. Pro residues) are colored white. In panels A, C, and D, key basic residues implicated in dsRNA binding, Arg208 and Lys221 are colored red. (B) The far UV CD spectrum of the NS1B-CTD*:16-bp dsRNA complex (orange line) is very similar to the sum of the CD spectra (blue line) of NS1B-CTD* and 16-bp dsRNA, indicating no major structural changes in either the protein or the dsRNA upon complex formation. (C) HADDOCK docking results showing a range of orientations for 16-bp dsRNA bound to NS1B-CTD, each of which is consistent with the NMR CSP data. (D) HADDOCK “hammerhead” model of the complex with best fit to the SAXS data, showing how sidechains of residues Arg208 and Lys221 bind the blunt end of the dsRNA. (E) Reciprocal space SAXS plot comparing experimental SAXS data (black) with SAXS scattering calculated for the best-fit blunt-end 5’OH-dsRNA binding “hammerhead” model (blue) and for best-fit two-state model comprised of a mixture of 23% 2:1 CTD : dsRNA dumbbell and 77% free dsRNA (red). For data in the range 0 < q < 0.25, the χ^2^‘s are 1.39 for the hammerhead model, and 0.99 for the two-state model. (F) Real space P(r) curves comparing experimental data (black) with curves calculated from the best-fit blunt-end hammerhead (blue) and two-state (red) binding models. (G) Schematic comparing the 1:1 hammerhead model (left) and the two-state binding models (right). The relatively high Porod-Debye exponent (2.7 units) indicates that the complex is flexible. See also **Supplementary Tables S2 and S3**, and **Figures S2, S3, and S7**.

Based on the three-dimensional structure of NS1B-CTD, we designed a construct with a single-site R238A mutation, referred to here as NS1B-CTD*, that suppresses oligomerization of NS1B without affecting the dsRNA binding activity of the CTD (Ma *et al*., 2016). NS1B-CTD* binds to 16-bp dsRNA with identical properties as the wild-type NS1B-CTD. However, these complexes also could not be crystallized. To characterize the three-dimensional structure of the NS1B-CTD* : 16-bp dsRNA complex we instead pursued an integrative structural biology approach, combining multiple biophysical measurements on these complexes in solution. First, far ultraviolet (UV) circular dichroism (CD) measurements were made for free NS1B-CTD*, free 16-bp dsRNA, and the stoichiometric complex formed between them. The sum of CD spectra (molar ellipticity) for NS1B-CTD* plus 16-bp dsRNA [(1:1) mixture] was compared with the molar ellipticity of the complex (**Figure 1B**). These data demonstrate that the complex formed between NS1B-CTD* and 16-bp dsRNA has CD spectral properties largely identical to the sum of the component CD spectra. In particular, the strong positive CD band with maximum at ∼ 260 nm, characteristic of A-form dsRNA (Clark et al., 1997; Chien et al., 2004), is identical for free and NS1B-CTD*-bound dsRNA (**Figure 1B**). As mentioned above, the backbone chemical shifts of NS1B-CTD* distant from the dsRNA-binding site were also not perturbed by complex formation (**Figure 1A, right**), indicating no overall structural changes to NS1B-CTD* protein upon complex formation. Taken together, these results demonstrate that complex formation does not significantly distort the structure of either the dsRNA or the NS1B-CTD* protein upon complex formation.

Structural modeling of the RNA-protein complex was next carried out using chemical-shift guided docking with the HADDOCK program (Dominguez et al., 2003; van Dijk and Bonvin, 2010; van Zundert et al., 2016), assuming a rigid docking process with no major changes in the structure of the RNA or protein upon complex formation. The individual NS1B-CTD (PDB 5DIL) (Ma *et al*., 2016) and 16-bp dsRNA (PDB 5KVJ) (**Supplementary Figure S1**) crystal structures were docked using the experimental chemical shift perturbation data (**Supplementary Table S2**) as outlined in detail in the Methods section. The resulting HADDOCK models of the complex (shown in **Figure 1C)** included both blunt-end and polyphosphate backbone (parallel) binding poses.

To provide additional structural data on the complex, small angle X-ray scattering (SAXS) experiments were next carried out. Guinier plot analysis of these SAXS data indicated little or no aggregation, low sample radiation damage, and appropriate P(r) curve shapes (**Supplementary Figure S2A and Supplementary Table S2**). SAXS curves that passed these quality checks were then compared against all of the models of the complex generated by HADDOCK using the FoXS server (Schneidman-Duhovny et al., 2016). The model that best fit the SAXS data (χ^2^ = 1.39) is shown in **Figure 1D**. In this best-fit single model, the 16-bp dsRNA interacts with the RNA-binding site on the surface of NS1B-CTD* in a blunt-end binding mode forming a “hammerhead” shape. In order to assess if these SAXS data distinguish blunt-end binding from a parallel backbone interaction like that observed for dsRNA binding to the N-terminal domain of NS1B (Yin *et al*., 2007), we also compared the P(r) curves for the best-fit (lowest χ^2^) blunt-end vs parallel binding models; the blunt-end binding (χ^2^ = 1.39) is a significantly better fit than the best-fit parallel binding modes (χ^2^ = 2.74) (**Supplementary Figure S3B**).

Although an excellent fit to the SAXS data was obtained with 1:1 NS1B-CTD : dsRNA “hammerhead” shaped complex (**Figure 1D, E**), careful analysis of the real space P(r) curves (**Figure 1F**) reveal the presence of a population of a larger-sized species. Indeed, electron density envelopes generated from the SAXS data also indicate a population of a dumbbell shaped object, consistent with a fraction of molecules in solution with blunt-end binding on both ends of the dsRNA (**Supplementary Figure S3A**). This is not unexpected since the two blunt ends of the 16-bp dsRNA have similar structures (**Supplementary Figure S1**). Fitting the SAXS data to a distribution of 23% dumbbell, < 1% hammerhead, and 77% free dsRNA provides an optimized fit to the data, with (χ^2^ = 0.99) and excellent fit agreement of the features of calculated and observe P(r) curves (**Figures 1E-G; Supplementary Figure S3**).

The best-fit blunt-end binding model for 16-bp dsRNA (**Figure 1D**) indicates a preferred binding mode, with NS1B-CTD binding at both ends of the simple dsRNA duplex substrate to from a dumbbell shape (**Figure 1G, two-state model**). Similar blunt-end binding (at one end of the dsRNA) is also observed in complexes formed between RIG-I and 5’ppp dsRNAs in the host innate immune response to viral infection (Jiang *et al*., 2011). This observation suggested the hypothesis that the blunt-end dsRNA binding mode of NS1B-CTD, like that of RIG-I, might be enhanced in 5’ppp-dsRNA molecules.

### NS1B-CTD preferentially binds blunt-ended 5’ppp-dsRNA, a pathogen-associated molecular pattern

Because the combined CD, NMR and SAXS data suggested that NS1B-CTD* binds to the RNA end, rather than to the RNA backbone, we next assayed whether CTD* has any RNA-end feature preferences. The SAXS data indicated binding could occur on either (or both) of the two exposed blunt ends of the 16-bp dsRNA substrate. In order to block binding at the end distal to our tested 5’ RNA modifications and ensure we were measuring affinity for our tested modifications, we employed hairpin (HP) RNA substrates. We generated three 10-base-pair 5’triphosphate (5’ppp) hairpin (HP) RNAs that were either blunt-ended (5’ppp ds10 HP RNA), had a 2-nt 3’ RNA overhang (5’ppp 3’ovg ds10 HP RNA), or a 2-nt 5’ RNA overhang (5’ppp 5’ovg ds10 HP RNA), to test whether NS1B-CTD* prefers blunt-ended or overhang RNA ends. We also tested an RNA substrate that lacked the 5’-triphosphate (5’OH ds10 HP RNA) to see whether CTD* had a 5’-triphosphate preference. Finally, we also generated a single strand, 10 nt RNA (5’ppp ss10) to test whether CTD* preferred double-stranded vs single-stranded RNAs. All RNAs had a 3’fluorescein moiety for use in fluorescence anisotropy or intensity assays (**Figure 2A**).

**Fig. 2.**
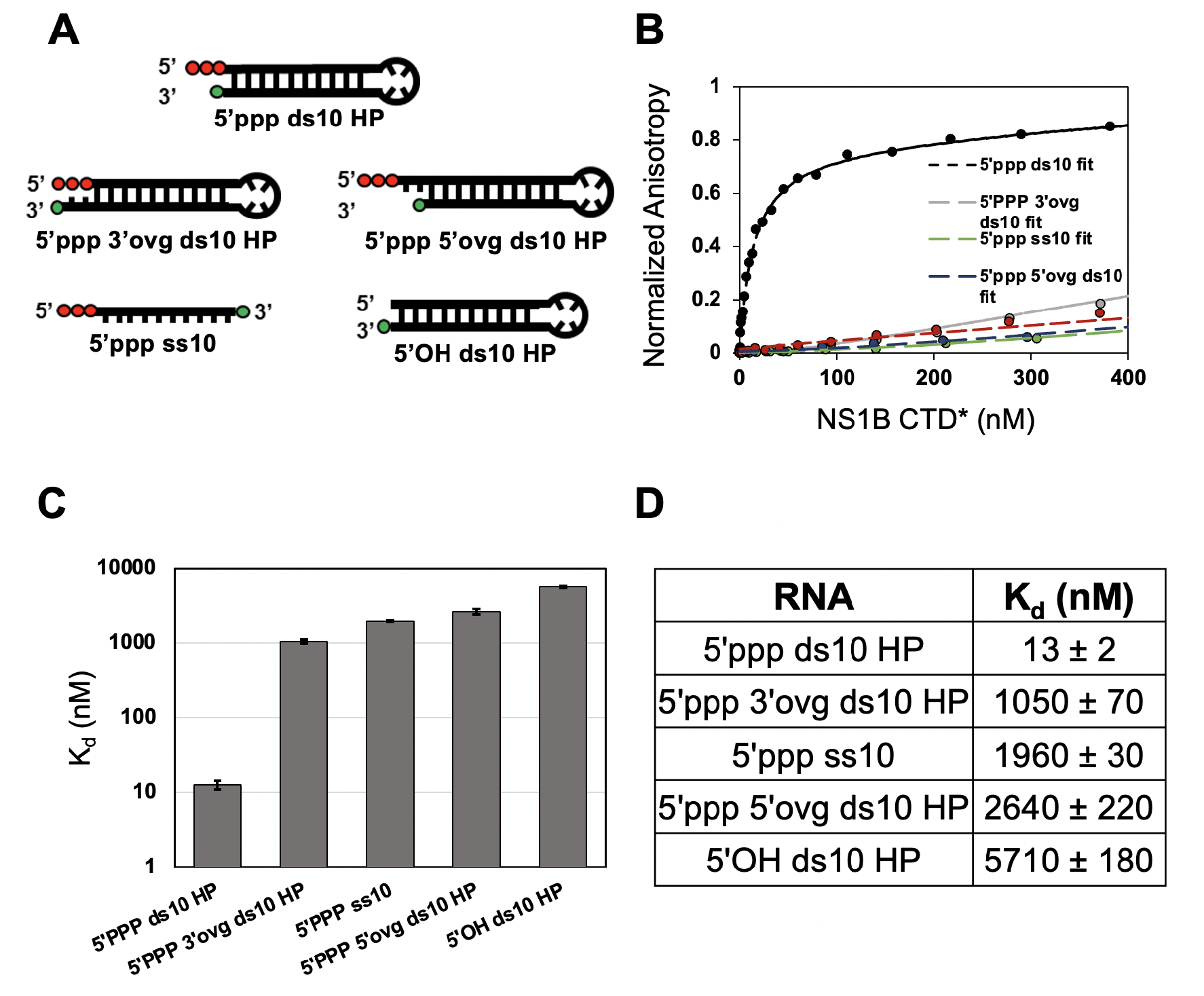
NS1B-CTD* has preference for blunt-ended 5’-triphosphate double-stranded RNA. (A) Schematics of the 10-bp hairpin RNAs with a 4-nt single-stranded loop used in this study. Red circles represent the 5’ppp moiety and green circle indicates the 3’-end fluorescein modification. Each overhang (ovg), whether at the 5’-end or the 3’-end, is 2-nt long. (B) Fluorescence anisotropy-based titration plots show the binding of various hairpin RNAs to NS1B-CTD*. Fluorescein-labeled hairpin RNA (20 nM) was titrated with increasing concentration of NS1B-CTD* and the resulting fluorescence anisotropy changes are shown for NS1B-CTD* concentrations between 0 and 400 nM. The anisotropy values are normalized from RNA alone (0) to maximum anisotropy (1) for each trial. Dashed lines indicate each trial’s regression fit to the binding equation. The 5’ppp ds10 HP RNA had at least two distinguishable binding phases (**Supplementary Figure S4**), and hence the data were fit to a double hyperbola. The K_d_ value of the tight binding phase is shown in (C) and (D) and the K_d_ of the weaker phase was 1800 ± 500 nM. 5’ppp 3’ovg ds10 HP RNA, 5’ppp ss10 RNA, and 5’ppp 5’ovg ds10 HP RNA showed cooperative binding character and were fit to the Hill equation. The Hill coefficient ranged from 1.2 to 1.4. 5’OH ds10 HP showed only one observable binding phase and was fit to a single hyperbola. Each binding experiment was repeated twice (n = 2). (C,D) The K_d_ values of the complexes are plotted in (C) and tabulated in (D). Errors are derived from errors of fits of two individual trials.

Out of all RNAs tested NS1B-CTD* bound tightly (K_d_ < 1 μM) only to 5’ppp ds10 HP RNA (**Figure 2B-D**), demonstrating a clear preference for blunt-ended, double-stranded RNAs with 5’triphosphate moieties. Significantly, NS1B-CTD* binds 440-fold more tightly to 5’ppp ds10 HP RNA (K_d_ = 13 ± 2 nM) than to 5’OH ds10 HP RNA (K_d_ = 5710 ± 180 nM). NS1B-CTD also has a significant preference (> 100-fold) for binding 5’ppp ds10 HP RNA over single-stranded 5’ppp ss10 RNA (K_d_ = 1960 ± 30 nM).

As protein titrations approached micromolar concentrations, we observed a second, weaker binding phase in our trials with 5’ppp ds10 HP RNA and NS1B-CTD* (**Supplemental Figure S4**) that could be delineated from the much tighter binding phase. We did not observe multiple binding phases for the rest of the tested RNA panel. Hence, it is possible that the K_d_’s we have reported for the rest of the panel are composite, apparent K_d_’s, where the two phases that resolved in 5’ppp ds10 HP RNA do not resolve clearly for the other tested RNAs. This would result in an apparent K_d_ that is weaker than the real K_d_ of NS1B-CTD* for a given RNA. However, this does not detract from the clear observation that blunt-ended, double-stranded, 5’ppp RNAs are significantly preferred relative to all other permutations tested here.

Finally, because the NS1B-CTD* construct contains a mutation, R238A, that suppresses NS1B-CTD homo-oligomerization (Ma *et al*., 2016), we also compared the binding of wild-type (WT) NS1B-CTD to 5’ppp ds10 HP RNA and to 5’OH ds10 HP RNA. We observed a similar preference for 5’ppp vs 5’OH (**Supplemental Figure S5A-B**), with the exception that the second binding phase showed a more significant increase in fluorescence anisotropy amplitude in WT NS1B-CTD compared to the R238A mutation, suggesting that the observed second phase may be related to CTD homo-oligomerization (**Supplemental Figure S5C**).

### Structural analysis the NS1B-CTD* : 5’ppp ds10 HP RNA complex

Considering the tight binding observed between NS1B-CTD* and 5’ppp ds10 HP RNA, the same set of NMR, CD, and SAXS experiments outlined above for the complex formed with 16-bp dsRNA were carried out for the NS1B-CTD* : 5’ppp ds10 HP complex. NMR resonance assignments for this complex were determined by conventional triple-resonance NMR methods, and backbone chemical shift perturbations (CSPs) relative to the free NS1B-CTD* were analyzed (**Supplementary Figure S6**). These CSPs plotted onto the X-ray crystal structure of NS1B-CTD* (**Figure 3A**) have a similar pattern to that observed for the complex with 16-bp 5’OH-dsRNA, with all the significant changes upon complex formation localized to the basic RNA binding face of the domain. Overall, the locations of CSPs on NS1B-CTD* due to RNA binding are similar for the 16-bp 5’OH-dsRNA and 5’ppp ds10 HP RNA substrates (**Supplementary Figure S7**. In particular, as with the 5’OH-dsRNA complex, no significant CSPs due to 5’ppp hairpin binding were observed on the opposite side of the domain, indicating little or no overall structural change upon complex formation.

**Fig. 3.**
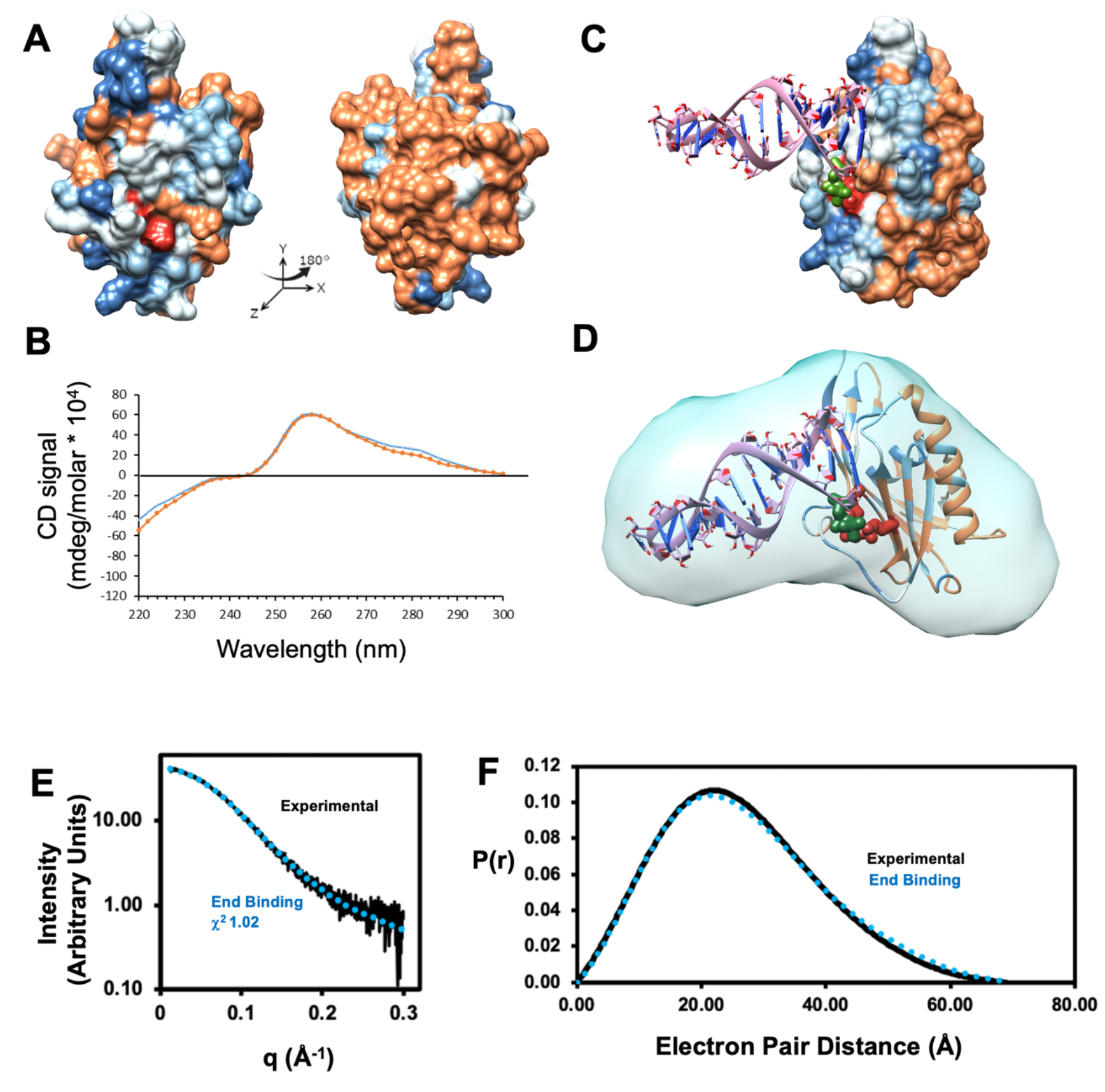
NS1B-CTD* binds 5’ppp ds10 HP in a blunt end orientation, with the triphosphate group binding into a basic a pocket. (A) CSPs from NS1B-CTD* bound to 5’ppp-ds10 HP RNA, mapped onto the surface of the NS1B-CTD* crystal structure. Surface residues are colored to indicate CSPs Δδ_ΝΗ_ > 95 ppb (dark blue), 95 > Δδ_ΝΗ_ > 65 ppb (light blue), Δδ_ΝΗ_ < 65 ppb (brown), or not determined (white). Note that in the lower salt buffer used in this RNA-binding study, the apo NS1B-CTD* aggregates to some degree, resulting in line broadening and larger cutoff values to define a significant CSPs (**Supplementary Figure S6**). In panels A,C, and D, key basic residues implicated in dsRNA binding, Arg208 and Lys221 are colored red, and the 5’ppp group of the hairpin RNA is colored green. (B) The far UV CD spectrum of the NS1B-CTD* : 5’ppp-ds10 HP RNA complex (orange line) is very similar to the sum of the CD spectra (blue line) of NS1B-CTD and 5’ppp-ds10 HP RNA, indicating no major structural changes in either the protein or the RNA upon complex. (C) HADDOCK model of the 5’ppp-ds10 HP RNA complex best fitting the SAXS data. (D) SAXS envelope of the best-fit NS1B-CTD* : 5’ppp-ds10 HP RNA complex, exhibiting a blunt end binding orientation, with side chains of basic residues Arg208 and Lys221 contacting the 5’ppp group. (E) Reciprocal space SAXS plot comparing experimental SAXS data (black points) with SAXS scattering calculated for the best-fit blunt-end 5’ppp-dsRNA binding model (blue line). For data in the range 0 < q < 0.25, the χ^2^ is 1.02. (E) Real space P(r) curves computed from the best-fit blunt-end model (blue) and experimental SAXS data (black). The relatively high Porod-Debye exponent (3.1 units) indicates that the complex is flexible. See also **Supplementary Tables S2 and S3**, and **Figures S2, S3C, S6, and S7**.

The UV CD spectra of free 5’ppp ds10 HP RNA, free NS1B-CTD*, and the complex formed between were next compared (**Figure 3B**). As observed for the complex with 16-bp 5’OH-dsRNA, the CD spectrum of the 5’ppp ds10 HP RNA complex is very similar to the sum of the CD spectra of 5’ppp ds10 HP RNA and NS1B-CTD* alone, indicating little or no change in the RNA conformation upon complex formation.

HADDOCK docking was then carried out using the NMR CSP data (**Supplementary Table S2**), as described in Methods. These modeling studies provided structural decoys with a wide range of interactions between NS1B-CTD* and bound 5’ppp ds10 HP molecule, including blunt-end and parallel binding, providing a range of models for subsequent SAXS analysis (**Supplemental Figure S3C)**. Next, SAXS studies were carried out as described in the Methods section. Guinier plot analysis indicated little or no aggregation, as well as consistency between expected and observed molecular weight, low sample radiation damage, and appropriate P(r) curve shapes (**Supplementary Figure S2B and Supplementary Table S3**). SAXS scattering curves were then compared to all models returned by HADDOCK using the FoXS server (Schneidman-Duhovny *et al*., 2016). The model with the best fit to the SAXS data, χ^2^ = 1.02, is shown in **Figures 3C and D**, together with a comparison of calculated and observed scattering (**Figure 3E, F).** In this model, the 5’ppp ds10 HP RNA interacts with the basic RNA-binding site on the surface of NS1B-CTD in a blunt-end binding mode, in which the 5’ppp group of the hairpin RNA is nearby to key basic residues Arg208 and Lys221, both of which are required for 5’ppp ds10 HP RNA binding. Improved data fits were not obtained by assuming various mixtures of free and bound complexes.

### NS1B-CTD competes with RIG-I for 5’ppp-dsRNA to suppress RIG-I activation

Tight blunt-end binding of NS1B-CTD* to 5’ppp-dsRNA, structurally resembling that of RIG-I to this same substrate, suggested that NSB1-CTD* would compete with RIG-I for 5’ppp dsRNA binding. RIG-I hydrolyzes ATP only while bound to RNA (**Figure 4A**). RIG-I’s ATPase is activated upon RNA binding and is essential for normal RIG-I function, and thus measuring the rate of ATP hydrolysis is a useful way to biochemically assay RIG-I activation (Devarkar *et al*., 2018). We titrated NS1B-CTD* against a fixed amount of RIG-I and 5’ppp ds10 HP RNA and measured the change in RIG-I ATPase rate. We observed that NS1B-CTD* inhibits RIG-I ATPase activity (**Figure 4B**). NS1B-CTD* inhibits 5’ppp-dsRNA activation of 15 nM RIG-I with an IC_50_ of 312 ± 32 nM.

**Fig. 4.**
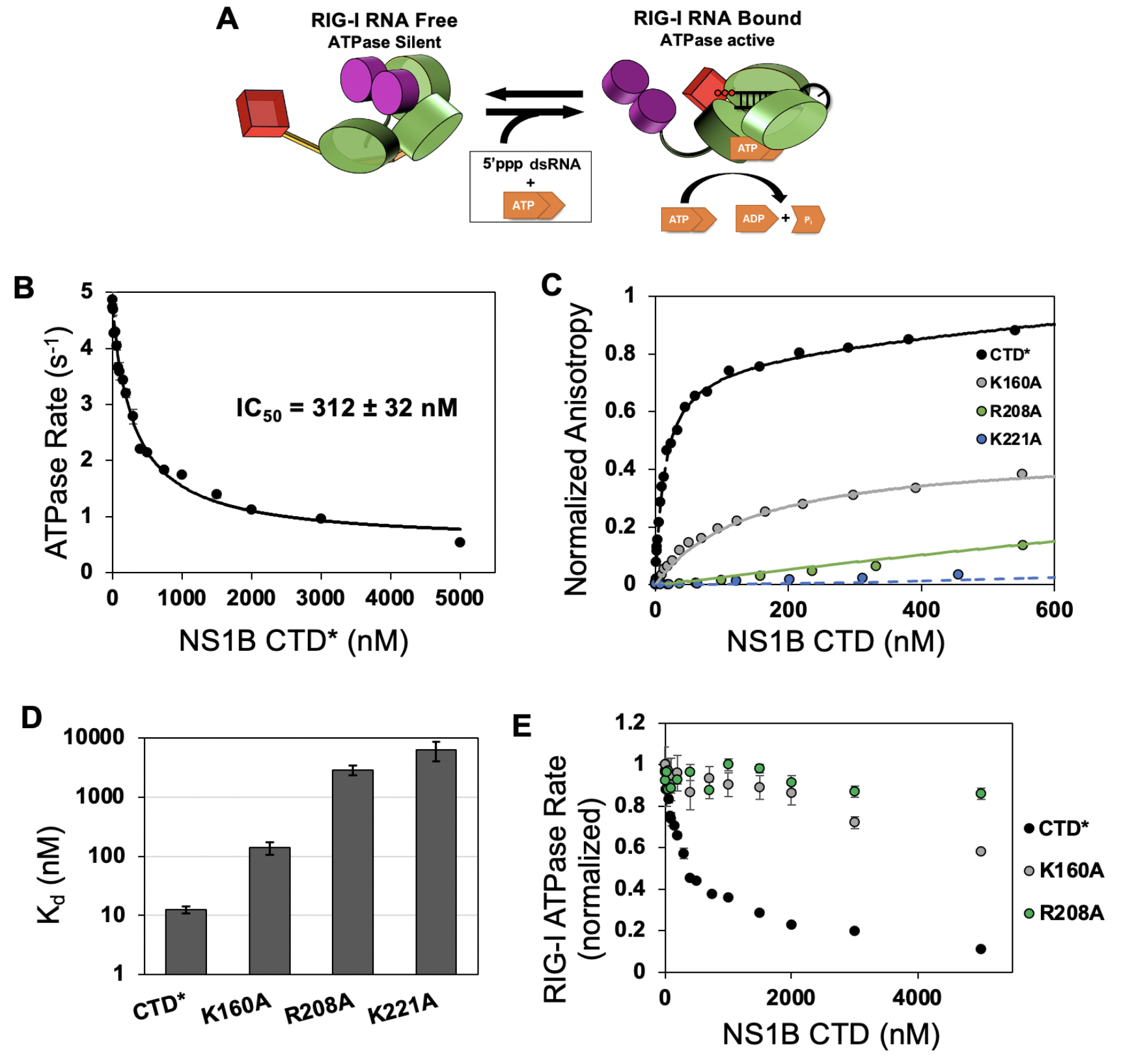
NS1B CTD* competitively inhibits RIG-I’s RNA-dependent ATPase activity. (A) Schematic of RIG-I’s RNA-dependent ATPase ability. RIG-I is broken into three main domains, the signaling CARDs (purple), the ATPase active helicase domain (green), and the 5’ RNA end binding domain RD (red). RNA free RIG-I is present in the autoinhibited state with undetectable ATPase activity. RIG-I is activated upon binding to dsRNA and hydrolyzes ATP to ADP and P_i_. Preventing its interaction with RNA would prevent RIG-I’s ATPase activity. (B) RIG-I (15 nM) was incubated with increasing concentration of NS1B-CTD* (0 – 5000 nM) and mixed with 5’ppp ds10 HP RNA (20 nM) and then ATP (2 mM) spiked with [γ-^32^P]ATP to start the reaction. The ATPase rate (M/N/s), which is M amount of P_i_ released per N amount of RIG-I per second) was obtained from a time course and plotted against NS1B CTD* concentration. The inhibition of RIG-I’s RNA-dependent ATPase activity was fit to a hyperbolic decay to calculate the IC_50_. Error bars are derived from each time course’s linear fit (n = 3). (C) Fluorescence anisotropy-based titrations show the binding of RNA-binding region mutants (Ma *et al*., 2016) of NS1B-CTD to 5’ppp ds10 HP RNA. The dashed lines are fit to a hyperbolic regression fit to calculate the K_d_. (D) The K_d_’s for each RNA-binding mutant to 5’ppp ds10 HP RNA. Error bars are derived from averaging two independent trials. (E) The RNA-dependent ATPase activity of RIG-I is inhibited to different extents by the RNA-binding mutants of NS1B-CTD*. The ATPase rates are normalized from 1 (no NS1B) to 0 (total inhibition).

### NS1B-CTD’s ability to inhibit RIG-I ATPase activity is dependent on its RNA-binding function

The NMR/SAXS model identified two basic residues in NS1B CTD that are predicted to be near the 5’-triphosphate (R208 and K221) in the complex. In addition, residue K160 is in the RNA-binding interface. These residues were shown to be important for 16-bp 5’OH-dsRNA binding (Ma *et al*., 2016). To verify that these basic residues are also required for 5’ppp hairpin RNA binding, we made alanine mutants at each of these sites and assayed their ability to bind 5’ppp ds10 HP RNA. We observed that each mutation severely impaired the binding of each mutant CTD to 5’ppp dsRNA (**Figure 4C,D**), with R208A and K221A, the two residues predicted to contact the 5’-triphosphate of the RNA, causing the most severe binding defects. We also assayed K160A and R208A NS1B-CTD* for their ability to inhibit RIG-I ATPase activity (**Figure 4E**). K160A showed an impaired ability to inhibit RIG-I ATPase activity, while R208A had no effect on RIG-I activation and therefore is unable to inhibit its activation by 5’ppp-dsRNA. These enzyme inhibition results correlate to the respective 5’ppp-dsRNA binding abilities of these mutants (*cf* **Figures 4C,D and E**). These data demonstrate that NS1B-CTD competes with RIG-I for target 5’ppp-dsRNA using the same region of its molecular surface shown to bind 5’ppp-dsRNA (**Figure 3A**).

### NS1B-CTD and not NTD guides preferential binding to 5’ppp-dsRNA in full-length NS1B

We next investigated whether NS1B-NTD, which has been previously been shown to bind along the RNA polyphosphate backbone of dsRNA in a parallel fashion (Yin *et al*., 2007), had any preference for 5’ppp versus 5’OH dsRNA. NS1B-NTD showed no significant difference in binding 5’ppp ds10 RNA compared to 5’OH ds10 HP RNA (**Figure 5A-B**) as measured by fluorescence anisotropy, indicating that NS1B-NTD does not prefer a 5’ppp moiety.

**Fig. 5.**
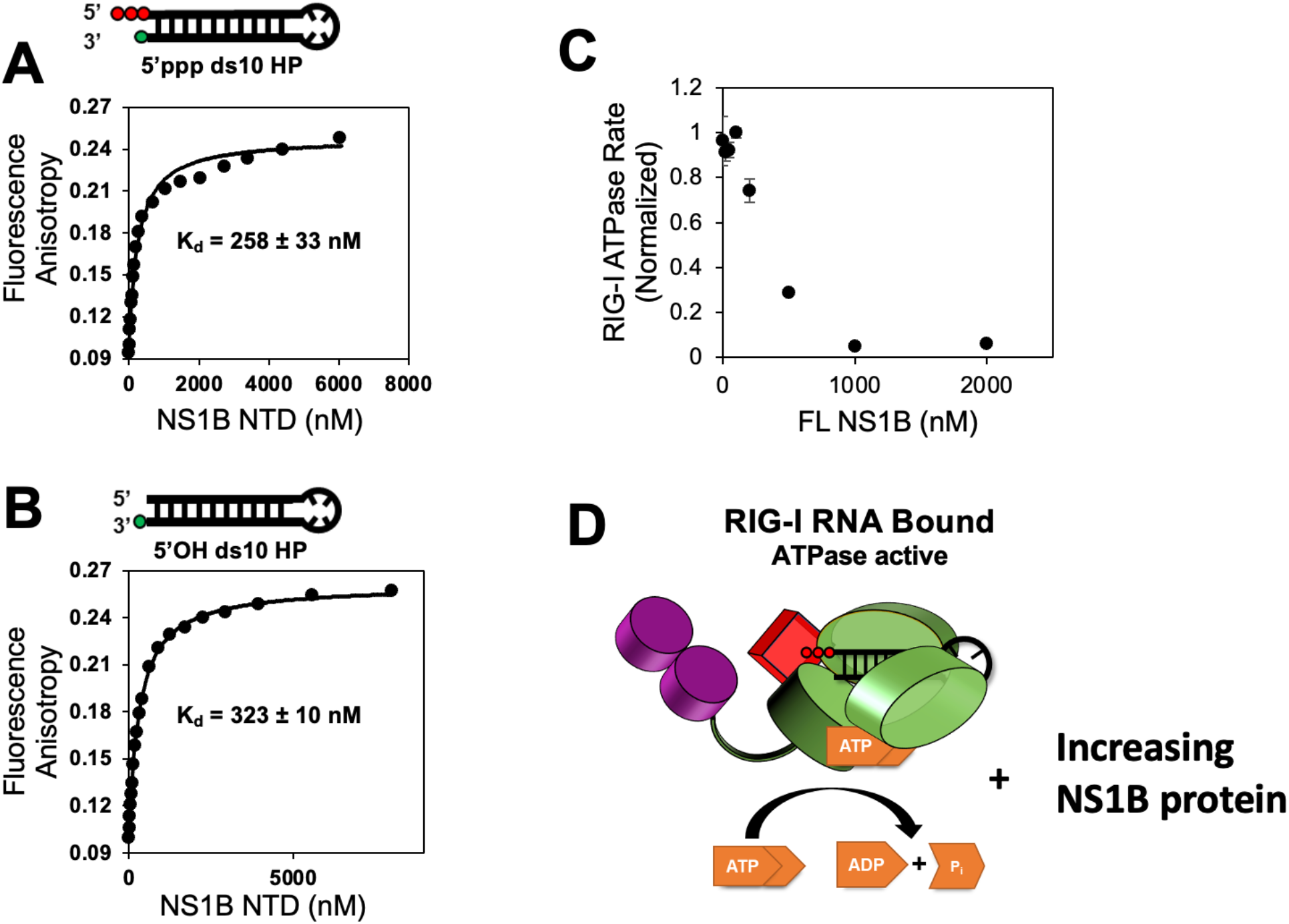
The NS1B full-length (FL) prefers 5’ppp over 5’OH RNA ends, and this preference is determined by its CTD, not NTD. (A) Fluorescein labeled 5’ppp ds10 HP RNA (20 nM) or (B) 5’OH ds10 HP RNA (20 nM) were titrated with increasing concentration of NS1B-NTD and the fluorescence anisotropy changes are shown. (C) RIG-I (20 nM) was incubated with increasing concentration of NS1B-FL (0 – 2000 nM) and mixed with 5’ppp ds70 RNA (20 nM) and then ATP (2 mM) spiked with [γ-^32^P]ATP to start the reaction. The normalized ATPase rate (M/N/s) obtained from the time courses of ATPase reactions are plotted against NS1B-FL concentrations. (D) Schematic diagram outlining the experimental design of panel C.

### Full-length NS1B suppresses RIG-I activation by 5’ppp-dsRNA

We also assessed whether full-length NS1B (FL-NS1B) is able to suppress 5’ppp-dsRNA activated RIG-I ATPase activity. In this study, we used a 5’ppp ds70 RNA substrate, an RNA that is sufficient to stimulate RIG-I signaling *in vivo* as compared to our hairpin RNA constructs (Devarkar *et al*., 2018). Like NS1B-CTD, the FL-NS1B also suppressed RIG-I activation by 5’ppp-dsRNA (**Figure 5C,D**). Thus, NS1B-CTD’s ability to target 5’ppp RNAs and inhibit activation of RIG-I is also observed for the full-length NS1B.

### NS1B-CTD RNA-binding activity inhibits activation of the RIG-I pathway in virus-infected cells

We next assessed our hypothesis, developed from these biochemical studies, that the 5’ppp-dsRNA binding function of the CTD of NS1B suppresses RIG-I activation in influenza B infected cells. PAMP binding to RIG-I in infected cells results in activation (phosphorylation) of the IRF-3 transcription factor that leads to the synthesis of interferon (Yoneyama *et al*., 2004; Yoneyama et al., 2005). We measured the levels of IRF-3 phosphorylation in cells infected with either wild-type virus or a mutant virus expressing an NS1B protein that is deficient in its CTD RNA-binding activity, specifically, a virus containing a R208A substitution in its NS1B protein that eliminates 5’ppp-dsRNA-binding activity. Infected cells were collected at 2, 4, 6, and 8 hours after infection, freeze-thawed three times and then total proteins were extracted using the PhosphoSafe reagent (Novagen) (**Figure 6A**). In cells infected with the R208A mutant virus, IRF3 was activated at all time points analyzed. This activation was not detected when infected cell extraction was carried out without the three freeze-thaws (Ma *et al*., 2016). In contrast, in wild-type virus infected cells the IRF3 activation detected initially at 2 hours after infection was suppressed at subsequent times after infection, presumably by newly synthesized wild-type NS1B proteins. These results demonstrate that a mutation that renders NS1B-CTD defective in 5’ppp-dsRNA binding function, and hence defective in competition with RIG-I for 5’ppp-dsRNA, impairs virus replication. NS1B-CTD inhibits RIG-I activation in virus-infected cells, as predicted by our biochemical studies, and its RNA binding activity is crucial for influenza B virus infection. To determine whether the N-terminal RNA-binding domain of NS1B, which binds dsRNA along the polyphosphate backbone (Yin *et al*., 2007), also inhibits IRF3, cells were infected with a mutant virus deficient in this RNA-binding activity; specifically, a virus with a R50A mutation in its NS1B protein. In cells infected by this mutant virus IRF3 activation was suppressed, indicating that the N-terminal RNA-binding site of the NS1B protein is not responsible for inhibiting IRF3 activation.

**Fig. 6.**
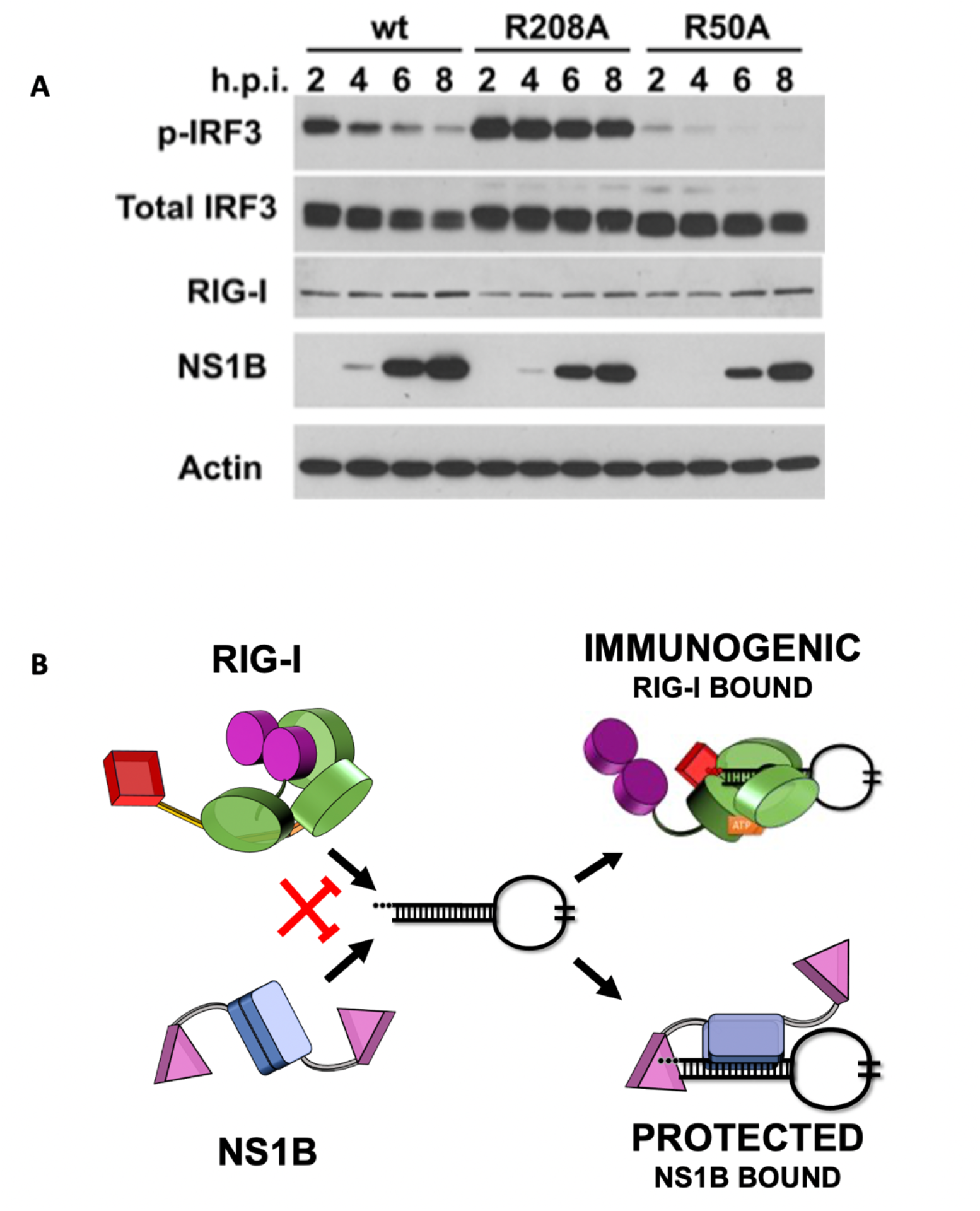
Proposed mechanism by which NS1B inhibits RIG-I activity. (A) Viral infection studies using WT and mutant influenza B viruses. Following infection with WT influenza B virus, or a mutant virus containing a R50A mutation in the N-terminal domain of NS1B, initial activation of IRF3 is suppressed as NS1 protein accumulates. However, in a virus containing a R208A mutation in the RNA-binding epitope of the C-terminal domain of NS1B, IRF3 activation is not blocked by accumulating NS1B protein. (B) NS1B exists as a NTD-mediated dimer in solution (blue rectangles). Both NS1B-CTD (pink triangles) and RIG-I recognize the 5’ppp-dsRNAs, such as the panhandle RNA region of influenza genomic vRNA, and compete for the 5’ppp-dsRNA blunt end. RIG-I binding vRNA culminates in interferon signaling, while NS1B binding sequesters otherwise immunogenic RNAs from RIG-I recognition. After NS1B binds, it can further oligomerize through repeated CTD and NTD dimerizations (Ma *et al*., 2016) potentially providing cooperativity and avidity; however, the role of such oligomerization is currently unknown.

## DISCUSSION

Each year influenza viruses cause highly contagious, respiratory disease that ranges from mild to severe. The detailed mechanisms by which influenza viruses evade the innate immune system are incompletely understood. In this study, we focus on influenza B and the structure and function of the recently characterized RNA-binding properties of the C-terminal domain of the IBV protein NS1B (Ma *et al*., 2016). Importantly, 5’ppp blunt-ended dsRNA is the primary PAMP of RIG-I (Yoneyama *et al*., 2005). Binding of 5’ppp blunt-ended dsRNA PAMPs by RIG-I causes structural rearrangement, activating its ATPase and helicase activities (Jiang *et al*., 2011), and initiating an innate immune response that includes phosphorylation and activation of the interferon regulatory factor 3 (IRF3) transcription factor. 5’ ppp blunt-ended dsRNA PAMPs also activate other RIG-I-like innate immune response receptors. 5’ppp single-stranded RNA (5’ppp-ssRNA) can also activate RIG-I at sufficiently high concentrations (Pichlmair et al., 2006).

The discovery of the 5’ppp specificity and role in immune evasion of NS1B extends our previous report that NS1B-CTD binds both single-stranded and double-stranded RNAs using a basic surface epitope that is conserved across IBV strains, but not present in the CTD of IAV NS1s (Ma *et al*., 2016). Mutation of each of the conserved basic residues in this surface epitope to alanine suppresses this RNA binding function, without disrupting the overall domain structure (Ma *et al*., 2016). The RNA-binding epitope of NS1B-CTD was mapped by NMR chemical shift perturbation studies of both ssRNA and dsRNA binding (Ma *et al*., 2016). Influenza B viruses genetically engineered with these mutations are also attenuated in cell-based viral replication assays (Ma *et al*., 2016). However, our previous study was not able to establish a clear relationship between this second RNA-binding function of NS1B and its function in viral infection.

The three-dimensional structure of NS1B-CTD bound to duplex 5’OH-dsRNA— determined here by combining NMR, X-ray crystallography, CD, single-site mutation, and SAXS data— unexpectedly revealed a blunt-end dsRNA binding mode. This blunt-end binding mode is structurally reminiscent of that observed in several RIG-I : dsRNA complexes (Lu *et al*., 2010; Jiang *et al*., 2011; Luo *et al*., 2012; Devarkar *et al*., 2016), suggesting that like RIG-I, the NS1B-CTD domain might prefer 5’ppp blunt-ended dsRNA substrates. Indeed, we observe an ∼ 440-fold enhancement in the affinity of NS1B-CTD for 5’ppp 10 HP vs 5’ OH 10 HP RNA. This binding to 5’ppp 10 HP RNA is attenuated by mutation of surface side chains in the previously characterized RNA-binding site of NS1B-CTD, including K160, R208, or K221, to alanine.

These mutations do not disrupt the overall structure of NS1B-CTD, based on NMR (^1^H-^15^N]-HSQC fingerprint spectra (Ma *et al*., 2016). Our SAXS / NMR studies demonstrate that the NS1B-CTD : 5’ppp 10 HP RNA complex does indeed utilize a blunt-end dsRNA binding mode analogous to that observed in 5’ppp dsRNA complexes with RIG-I. Full-length NS1B, guided by its CTD, can compete with RIG-I for 5’ ppp dsRNAs, thus suppressing RIG-I signaling activation both in biophysical studies and in virus-infected cells. These observations suggest that the 5’ppp blunt-ended dsRNA binding function of NS1B may have evolved to suppress the activation of the host antiviral response by 5’ppp dsRNA blunt ends of vRNAs and viral replication intermediates that otherwise would activate RIG-I and the innate immune response to infection.

In both influenza A and B viruses, the 3’ and 5’ ends of each RNA segment are complementary, forming a 5’ triphosphorylated dsRNA “panhandle” (Desselberger *et al*., 1980; Hsu *et al*., 1987; Cheong *et al*., 1999; Bae *et al*., 2001; Hofacker et al., 2004). 5’-triphosphorylated dsRNAs act as PAMPs for activation of RIG-I. RIG-I can also be activated by 5’ppp-ssRNAs. Most of our understanding of how influenza viruses suppresses this pathway comes from studies of IAV. Pichlmaier et al. have reported that in IAV infection RIG-I is activated by viral genomic 5’ppp-ssRNA, and that this activation is blocked by NS1A, which was found in a complex with RIG-I in infected cells (Pichlmair *et al*., 2006). However, these studies did not identify how NS1A blocked RIG-I activation, and could not exclude that the activation of RIG-I is due to hairpin formation by 5’ppp-ssRNAs. More recently, Liu et. al (Liu *et al*., 2015) have shown that the IAV panhandle structure is directly involved in RIG-I activation and interferon induction. In porcine alveolar macrophages the viral genomic coding region (i.e. the “pan’) is dispensible for RIG-I-dependent IFN induction, while a synthetic IAV panhandle RNA was found to directly bind to RIG-I and stimulate IFN production (Liu *et al*., 2015). Although there are not yet similar reports for IBV genomic panhandles activating RIG-I, IAV and IBV panhandles have similar 5’ppp hairpin structures (Desselberger *et al*., 1980; Hsu *et al*., 1987; Hofacker *et al*., 2004). vRNAs resulting from IBV replication, and also those present in the cell before packaging, as well as vRNA replication defective intermediates with 5’ppp blunt-ended dsRNA structures (Baum and Garcia-Sastre, 2011), may also function as PAMPs for RIG-I that are suppressed by NS1B.

IBV mutant viruses with alanine substitutions of key NS1B residues R156A, R160A, and R208A are severely attenuated in infecting A549 cells (Ma *et al*., 2016). In this same study, an IBV virus with the R221A-NS1B mutation was so severely impacted that it could not be grown in culture at all. Significantly, in the NMR/SAXS model of the complex formed between NS1B-CTD and 5’ppp ds10 HP RNA, all four of these basic residues are located in the interface between the RNA-binding epitope of NS1B-CTD and the blunt end of 5’ppp ds10 HP RNA.

These combined observations lead to the hypothesis that at least one cellular function of the 5’ppp dsRNA-binding activity of NS1B is to suppress the host antiviral response by competing with RIG-I (and/or RIG-I - like innate immune receptors) for various 5’ppp PAMPs that result from viral infection (as illustrated in **Figure 6B**). To test this hypothesis, cell-based viral infection studies were carried out with a mutant influenza B virus. Infection of A549 cells with WT IBV virus results in initial activation of the host antiviral response, including activation of the RIG-I pathway and subsequent phosphorylation of transcription factor IRF3, leading to enhanced expression of type-I interferons and other response genes. However, rising levels of NS1B during the viral infection process then attenuate RIG-I pathway activation, suppressing the innate immune response, and allowing successful viral infection. Infection with a mutant [R208A]-NS1B influenza B virus results in initial RIG-I pathway activation. However, because [R208A]-NS1B has impaired 5’ ppp dsRNA binding ability, it cannot compete with RIG-I for 5’ppp dsRNAs, such as the genomic panhandle of IBV, and suppress this innate immune response. This mutation results in a virus impaired in innate immune evasion, leading to sustained RIG-I pathway activation and innate immune signaling during infection, and hence attenuating viral infection.

The novel mechanism described here by which the NS1B protein suppresses the interferon response differs from two mechanisms that are employed by the NS1A protein of influenza A virus. In one, the NS1A protein binds and sequesters the 30 kDa subunit of the cleavage and polyadenylation specificity factor (CPSF), the cellular factor that is required for the 3’-end processing of cellular pre-mRNAs (Nemeroff et al., 1998).

This interaction inhibits the 3’ processing of cellular pre-mRNAs and hence the production of mature mRNAs, including mature IFN and other antiviral cellular mRNAs (Das et al., 2008). In a second proposed mechanism, the NS1A protein suppresses the RIG-I signaling pathway by binding host factor TRIM25, thereby interfering with the correct positioning of the PRYSPRY domain of TRIM25 that is required for substrate ubiquitination (Koliopoulos et al., 2018). Interestingly, only one of these mechanisms is sufficient for some circulating influenza A viruses that do not activate the RIG-I pathway (Kuo et al., 2010; Krug and Garcia-Sastre, 2013). Influenza B virus does not appear to use either of these mechanisms, as NS1B has not been reported to bind either CPSF or TRIM25.

Similar to our proposed model of NS1B competing with RIG-I for 5’ppp dsRNA ends, Ebola viruses (EBOV) employ an analogous strategy to suppress RIG-I activation and inhibit alpha/beta interferon production (Cardenas et al., 2006; Leung et al., 2010; Bale et al., 2013). Isothermal titration calorimetry studies demonstrated ∼ 15 fold tighter binding by Ebola VP35 interferon inhibitory domain IID by in vitro transcribed 8-bp 5’ppp dsRNA compared to the same substrate with 5’OH (Leung *et al*., 2010). The virally-encoded VP35 domain IIB structurally caps dsRNA using an asymmetric dimer, where one molecule specifically binds the RNA blunt end and another primarily interacts with the RNA backbone (Leung *et al*., 2010), with features analogous to those proposed here for full-length influenza NS1B binding 5’ppp-dsRNA (**Figure 6B**). This blunt-end-targeted binding may nucleate VP35 coating of the RNA backbone, protecting the RNA from RIG-I recognition (Leung *et al*., 2010). Despite the analogy between this mechanism and the one described here for IBV NS1B competing with RIG-I for 5’ppp viral PAMPS, there is no significant structural or sequence similarity between these EV and IBV structural proteins.

The genomic RNAs of several other negative-strand viruses, including rabies, bunya, and measles viruses, also have partially complementary sequences in their 5’ and 3’ regions which can activate RIG-I (Hofacker *et al*., 2004; Fujita, 2009; Schlee *et al*., 2009; Liu *et al*., 2015). For example, a synthetic rabies viral panhandle (without a pan) has been reported to activate RIG-I (Schlee *et al*., 2009). Although IAV genomic RNAs can form 5’ppp panhandles, the CTD of NS1 from Udorn influenza A virus does not bind 5’OH dsRNA (Ma *et al*., 2016) or 5’ppp ds10 HP RNA (data not shown), and lacks the key conserved basic sites that mediate blunt-end binding by the CTD of NS1 from influenza B viruses (Ma *et al*., 2016). It remains to be seen if the 5’ppp-dsRNA binding mechanism used by the NS1B protein of IVB is unique to this viral lineage, or if some of these other negative-strand viruses also encode non-structural proteins that competitively suppress RIG-I activation by blunt-end binding to 5’ppp-dsRNA PAMPs.

In conclusion, we have identified a novel and unanticipated role in viral infection for NS1B and for the RNA binding function of its CTD: i.e. sequestering 5’ppp dsRNAs that is otherwise recognized by RIG-I and RIG-I -like cellular innate immune receptors. This function of NS1 allows IBV to productively replicate without triggering a strong host antiviral response. Unlike the NS1B-NTD which binds nonspecifically to the dsRNA backbone, NS1B-CTD preferentially binds 5’-ppp, blunt-ended, double-stranded RNA motifs, like those associated with PAMPs of RIG-I-like innate immune receptors. These PAMPs include the NS1B genomic panhandle and viral replication intermediates. Hence, specific inhibitors of this 5’ppp dsRNA binding function of NS1B could potentially be developed as influenza antivirals. These results reveal a novel mechanism of innate immune evasion by influenza B virus and open the door to new approaches for antiviral drug discovery.

## ACKNOWLEDGEMENTS

We thank Profs. Eddy Arnold, Joseph Marcotrigiano, Vikas Nanda, and Monica Roth for helpful discussions and intellectual input. We also thank the SIBYLS beamline staff for expert advice and assistance, Dr. Richard Gillilan and MacCHESS for onsite training, Dr. Jesse Hopkins and the BioCAT facility for experimental assistance. We also thank Viviane Liao for assistance. This work was supported by National Institute of Health Grants R35-GM141818 (to G.T.M.), R35-GM118086 (to S.S.P.), R01-GM120574 (to G.T.M), R21S-AI117510 (to G.T.M., R.W.), R01-AI11772 (to R.M.K.), P01-CA092584 (to J.A.T., S.T., G.H.), R35-CA220430 (to J.A.T.), DOE BRAVE Taskforce 5 Program (to J.A.T., S.T., G.H.), Robert A. Welch Chemistry Chair (G-0010 to J.A.T.), and the RPI Constellation Endowed Chair fund (to G.T.M.).

## METHODS

## KEY RESOURCES TABLE

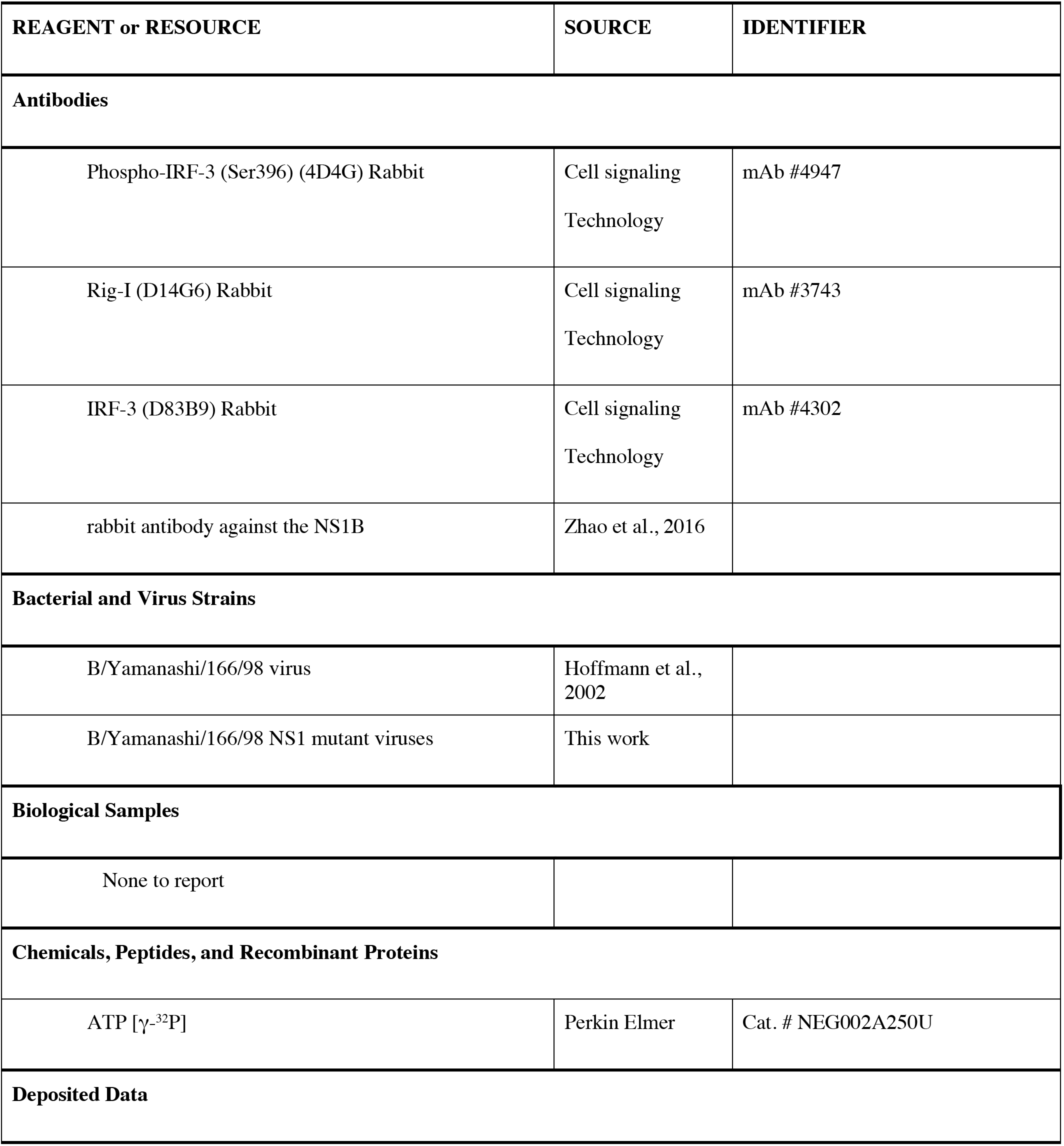
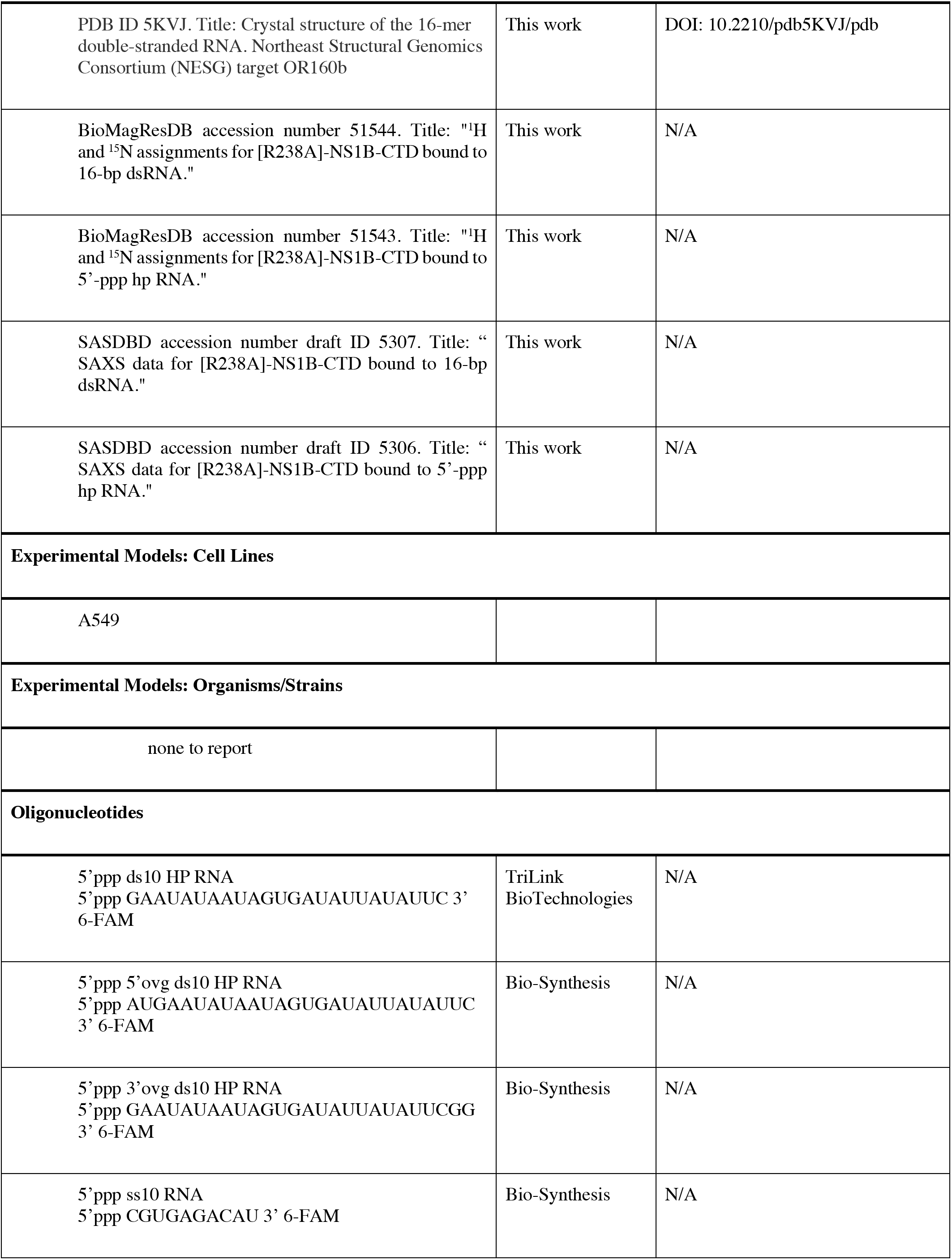
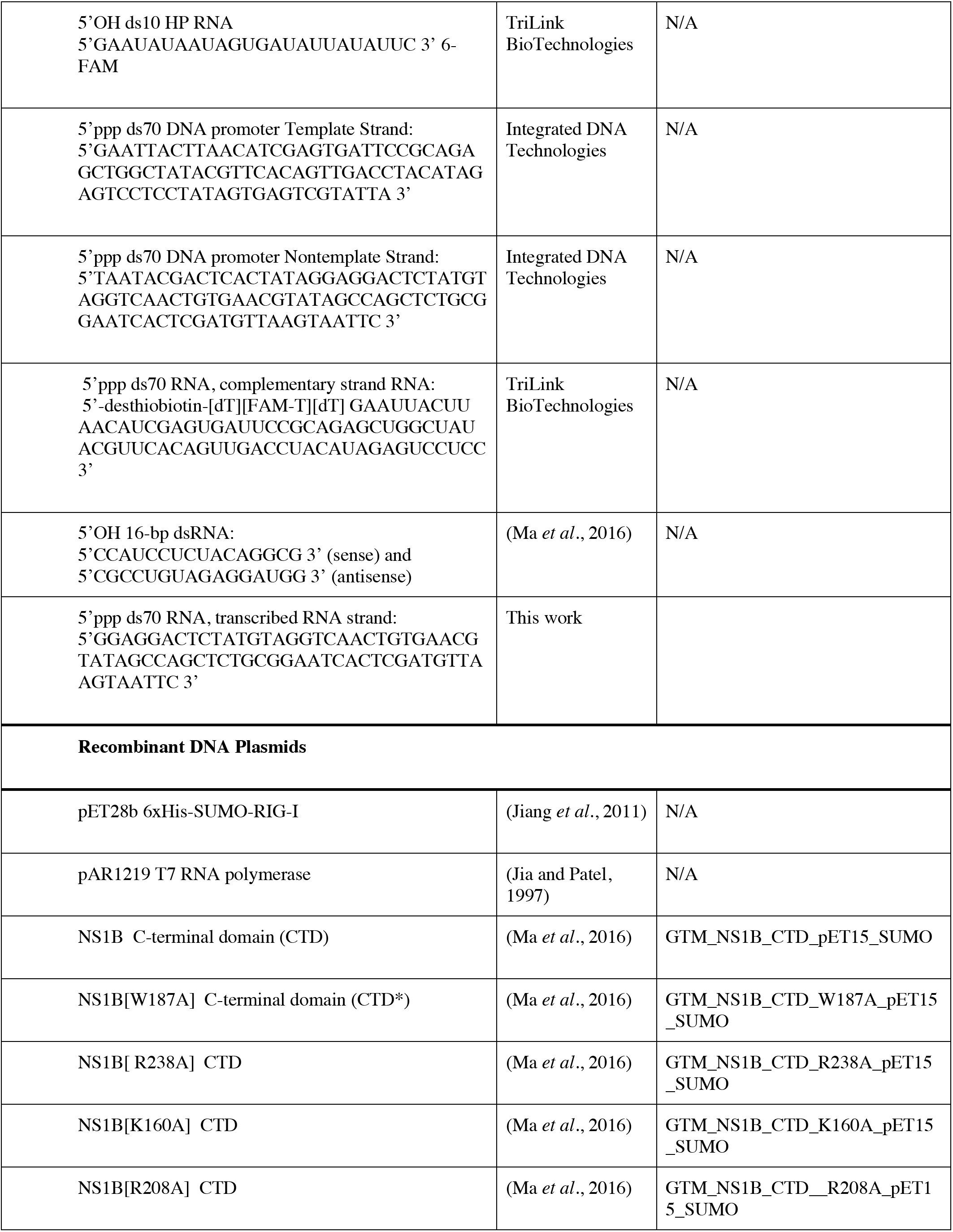
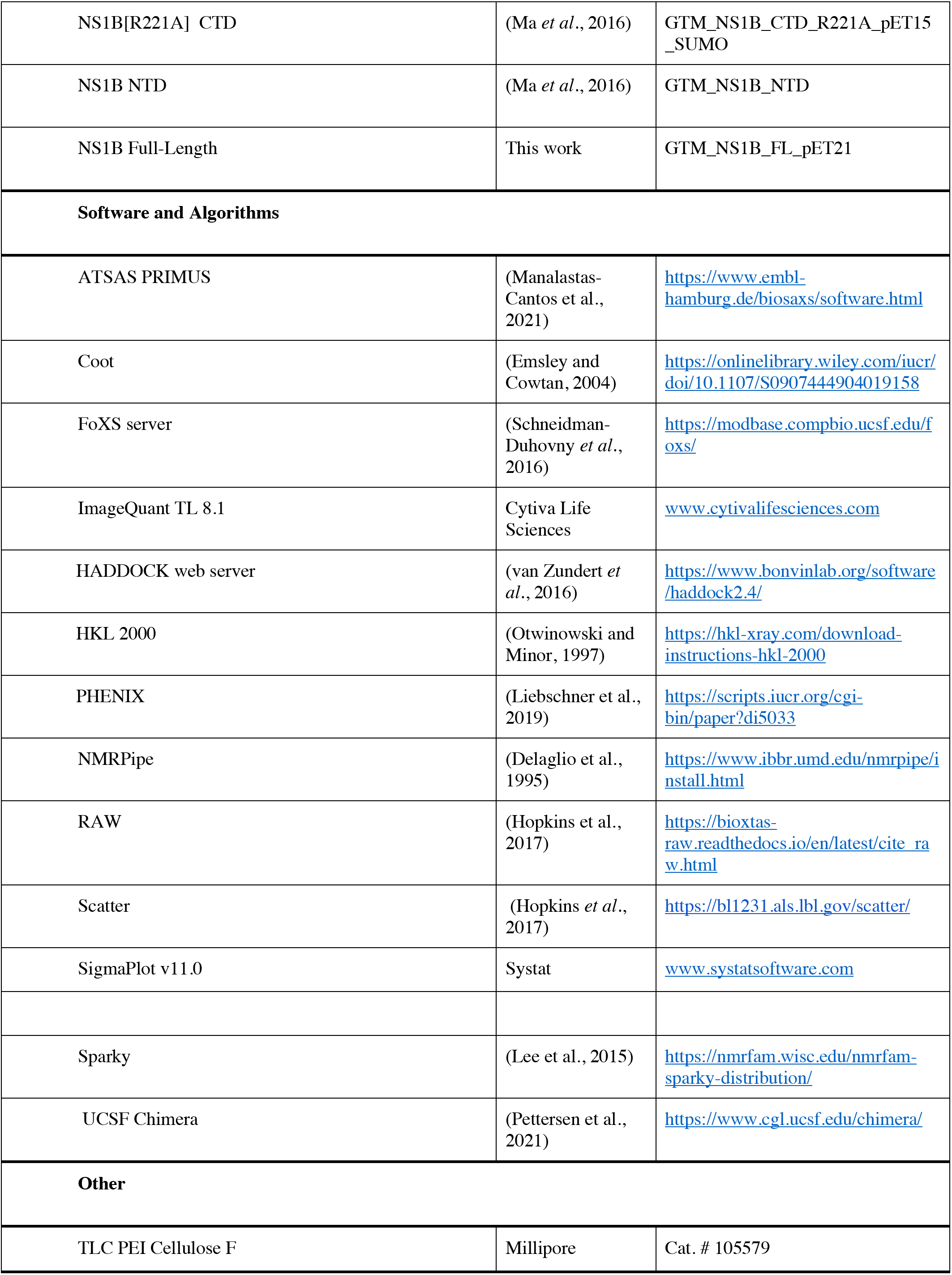

## RESOURCE AVAILABILITY

### Lead contact

Further information and requests for resources and reagents should be directed to and will be fulfilled by the designated contacts G. T. Montelione and S. Patel.

## MATERIALS AVAILABILITY

All materials generated in this study are available upon request.

## DATA AND CODE AVAILABILITY

The following data sets generated during this study have been deposited in public data repositories:

1. Protein Data Bank PDB id 5KVJ. Title: “Crystal structure of a 16-mer double-stranded RNA substrate that binds to influenza NS1”. DOI: 10.2210/pdb5KVJ/pdb
2. BioMagResDB accession number: 51543: Title: “^1^H and ^15^N assignments for [R238A]-NS1B-CTD bound to 5’-ppp hp RNA.”
3. BioMagResDB accession number: 51544: Title: “^1^H and ^15^N assignments for [R238A]-NS1B-CTD bound to 16-bp dsRNA.”
4. SASDBD accession number draft ID 5307. Title: “SAXS data for [R238A]-NS1B-CTD bound to 16-bp dsRNA.”
5. SASDBD accession number draft ID 5306. Title: “SAXS data for [R238A]-NS1B-CTD bound to 5’-ppp hp RNA.”

## METHODS DETAILS

### Protein expression and purification: NS1B-CTD and mutants

NS1B-CTD (residues 141-281 of strain B/Lee) and NS1B-CTD* (with W187A single-site mutation) were prepared as described in previously (Ma *et al*., 2016) using plasmids GTM_NS1B_CTD_ pET15_SUMO and GTM_NS1B_CTD_W187A_pET15_SUMO, respectively, with ampicillin resistance. Other constructs of single-site mutants of NS1B-CTD were prepared as described previously (Ma *et al*., 2016), using the expression plasmids listed in the STAR Methods Table. These provide a His_6_-SUMO N-terminal fusion with the targeted NS1B-CTD constructs. Plasmids were transformed into *E. coli* BL21(DE3) Mgk cells and fermented in standard LB or MJ9 minimal isotope-enrichment (Jansson et al., 1996) media, to produce unlabeled and isotope-enriched proteins respectively. MJ9 media contained (^15^NH_4_)_2_SO_4_ and/or D- [U^13^C]- glucose as the soles of nitrogen and carbon, respectively. Cells were first grown at 37 °C to an absorbance of ∼ 0.6 at 600 nm, and then expression was induced by adding isopropyl-β-D-thiogalactoside (IPTG) at a concentration of 1 mM and incubating at 17 °C for ∼ 18 hrs. The cells were then pelleted by centrifugation at 5,000 x g for 45 mins and the pellet was either prepared directly for purification or stored at -80 °C.

Purification buffers were freshly prepared before each purification. Ionic strength plays an important role in protein-RNA interactions, and special attention to the optimized buffer and salt concentrations is essential for replicable experiments. Ten buffers summarized in **Supplementary Table S4** were used for protein production and structural characterization: (i) Ni binding buffer (buffer BB), (ii) Ni elution buffer (buffer EB), (iii) Low Salt buffer (buffer LS), (iv) Heparin buffer A (buffer A), (iv) Heparin buffer B (buffer B), (vi) NS1B-CTD buffer (buffer 2K), (vii) NS1B-CTD RNA binding buffer (buffer 1K), (viii) NS1B-full length buffer (buffer FL), (ix) RNA Binding Buffer, and (x) Storage Buffer. All buffers were filtered with a 0.1 µm filter to sterilize. Buffers used for SAXS studies were double-filtered using an additional 0.02 µm filter. to ensure removal of small particles which can seep through the 0.1 µm filter.

Pellets for cell lysis were suspended in Buffer BB, transferred into metal cups, the cups were floated in an ice bath, and the samples were sonicated with 10 sonic pulse cycles of 30 seconds on / 30 seconds off. The resulting protein slurry was passed through a 0.1 µm filter, and applied to a Ni-NTA column on an AKTAExpress chromatography system, using a flow rate of 0.5 mL / min. The column had a loading step followed by 2X volume washing step, and finally an elution step. The loading and washing steps were done with the buffer BB, while the elution was done with buffer EB. There was no gradient transition. The fractions with a high UV absorbance (read from the chromatogram) were analyzed by SDS-PAGE. All relevant fractions were pooled together, adjusted to 5 mM DTT, and Ulp1 SUMO protease with an N-terminal hexaHis tag [prepared as described in Mazzei et al (Mazzei et al., 2023)] was added at a 20 : 1 or 50 : 1 protein: protease ratios to cleave the SUMO tag from the SUMO-NS1B-CTD fusion. SDS-PAGE was used to assess cleavage. Once > 90% cleavage was observed, the sample was run on a heparin column to remove bound nucleic acids, using an AKTAPure FPLC chromatography system. The heparin column was run by loading the sample in Heparin buffer A, washing the column with buffer A, and then increasing salt in a gradient up to 1 M NaCl by applying a gradient of buffer B over a volume of ∼45 mL. SDS-PAGE was used to identify the fractions containing NS1B-CTD and to assess purity of the eluted protein. It is important to note that NS1B-CTD* has no Trp residues, and the UV absorbance at 280 nm for the protein with no tag is very low due to its low molar extinction coefficient at this wavelength. The samples containing NS1B-CTD were then pooled together and loaded directly back into the Ni column using the same protocol described above. In this step, care was taken to collect all samples containing the cleaved NS1B-CTD, including flow through and wash steps, as the protein no longer has a 6XHis-SUMO tag and will not bind to the column. This last step is done to remove any traces of uncleaved fusion protein and the His_6_-tagged SUMO protease. Note that in some cases, multiple passes through the Ni-NTA column were needed to remove all of the uncleaved His_6_-SUMO-NS1B-CTD fusion protein. The final samples were validated by MALDI-TOF mass spectrometry, and assessed to be > 98% homogeneous by SDS-PAGE with Coomassie blue staining.

The resulting purified NS1B-CTD, NS1B-CTD*, and other single-site constructs prepared as described above are very sensitive to buffer conditions and have a tendency to aggregate. Aggregation causes light scattering, which was assessed by UV spectroscopy, paying special attention to the 260 nm / 280 nm ratio, and confirming that the apparent absorbance spectrum goes to zero before reaching 300 nm. We observed samples that have a long apparent absorbance tail that extends beyond 300 nm due to aggregation. When samples contained this tail due to light scattering, the sample was centrifuged at 8,500 x g for 10-20 minutes. The supernatant was then transferred to another tube, being careful not to mix the sample or disturb the pellet, which is nearly invisible to the human eye. The UV spectrum is taken again and compared to the spectrum previous to centrifugation. This is repeated until the sample provides a suitable UV absorbance spectrum indicating little or no light scattering above 300 nm due to aggregation, and a A_260_ / A_280_ ratio < 0.7. The sample was then put on ice and the concentration determined by absorbance at 280 nm, using the extinction coefficient estimated from the protein sequence, 1490 Abs Units/(M-cm) at 280 nm.

### Protein expression and purification: full-length NS1B

Full-length (FL) NS1B from strain B/Lee was cloned into vector pET21_NESG with ampicillin resistance to create plasmid GTM_NS1B_FL_pET21, encoding the full-length NS1B protein with a C-terminal hexaHis tag. Full-length NS1B was produced by fermentation, Ni-NTA purifications, and preparative gel filtration as described elsewhere (Xiao et al., 2010; Acton et al., 2011), and then buffer exchanged into FL buffer for storage.

### Protein expression and purification: RIG-I and T7 RNA polymerase

Human RIG-I was expressed using pET28 SUMO vector in *Escherichia coli* strain Rosetta (DE3) (Novagen). The cell lysate soluble fraction was purified using a Ni^2+^-nitrilotriacetate (QIAGEN) column, followed by Ulp1 SUMO protease digestion to remove 6xHis-SUMO tag. It was further purified by hydroxyapatite (CHT-II, Bio-Rad) and heparin Sepharose column chromatography (GE Healthcare). Purified protein was dialyzed into 50 mM HEPES pH 7.5, 50 mM NaCl, 5mM MgCl_2_, 5 mM DTT, and 10% glycerol overnight at 4°C, then snap frozen in liquid nitrogen and stored at -80°C (Jiang *et al*., 2011).

T7 RNA polymerase was expressed using pAR1219 vector in *Escherichia coli* strain BL21 (Novagen). Cell lysate was purified in three steps: SP-Sephadex (Sigma Aldrich), CM-Sephadex (Sigma Aldrich), and then DEAE-Sephacel (Sigma Aldrich). Finally, purified protein was dialyzed into Storage Buffer (20 mM sodium phosphate pH 7.7, 1 mM Na_3_-EDTA, 1 mM dithiothreitol, 100 mM NaCl, and 50% (v/v) glycerol) (Jia and Patel, 1997).

### RNAs sample preparation for binding affinity studies

RNAs for fluorescence-based binding affinity measurements were chemically synthesized and HPLC purified by either TriLink BioTechnologies. Inc. or Bio-Synthesis, Inc. Purity was confirmed using mass spectrometry and HPLC. Lyophilized RNA was resuspended in 50 mM Tris-Cl, pH 7.4, and 20 mM NaCl. Concentrations of RNAs were determined by measuring their absorbance using a NanoDrop spectrophotometer at 260 nm in 7 M guanidium HCl, using each RNA’s predicted extinction coefficient.

The 70-mer single-stranded RNA containing the 5’ppp was *in vitro* transcribed using T7 RNA polymerase. The promoter DNA (2 µM) was incubated with 2 mM spermidine (Thermo Fisher Scientific); 5 mM of each ATP, GTP, UTP, and CTP (Thermo Fisher Scientific); 5 µg inorganic pyrophosphatase (New England BioLabs); 50 U RNasin (Promega), and 1 µM T7 RNAP in a buffer containing 250 mM HEPES pH 7.5, 30 mM MgCl_2_, and 40 mM DTT at 30 °C for 3 hours. The reaction was quenched with addition of 75 mM EDTA, run through a preparative 15% polyacrylamide, 6 M urea gel with a 1x TBE running buffer. The band corresponding to the transcribed RNA product was then cut from the gel and further purified using gel electroelution at 150V, taking fractions every two hours, and then run overnight at 20V, all in 1x TBE running buffer. Collected fractions were then concentrated, and the sample was sent for mass spectroscopy analysis to confirm purity. Both the transcribed triphosphate strand and chemically synthesized complementary strand were annealed at a 1:1 ratio in 50 mM Tris-Cl, pH 7.4, and 20 mM NaCl. The extinction coefficient of the transcribed RNA was calculated using a nearest neighbor molar extinction calculator (idtdna.com).

### RIG-I ATPase inhibition assays

NS1B’s ability to inhibit RIG-I ATPase activity was measured at constant RIG-I (15 nM) and RNA (20 nM), and increasing NS1B or NS1B-CTD construct concentration (5 nM – 5 µM), in the presence of 2 mM ATP (labeled with [γ-^32^P] ATP). A time course (0-45 minutes) of the ATPase reactions were performed in RNA Binding Buffer (50 mM MOPS pH 7.4, 5 mM DTT, 5 mM MgCl_2_, 0.01% Tween20) at 25 °C. The time course was measured at 0, 15, 30, and 45 minutes to generate three independent hydrolysis rates per NS1B concentration (n = 3). RIG-I and NS1B constructs were mixed before adding RNA, so as to not bias the inhibition assay with RNA prebinding to either RIG-I or NS1B constructs. Reactions were stopped using 4 N formic acid and analyzed with PEI-Cellulose-F TLC (Millipore) run with 0.4 M potassium phosphate buffer (pH 3.4). TLC plates were exposed to a phosphoimaging plate, visualized using a Typhoon phosphor-imager, and then quantified using ImageQuant software. ATPase rates at each concentration of NS1B construct were determined by plotting the molar [P_i_] against time (s), and these rates were then plotted as a function of RNA concentration. When quantified, these graphs were fit with a hyperbolic decay equation (Eqn. 1) to determine maximal ATPase rate (*k*_atpase_) and estimate the IC_50_ of NS1B constructs.

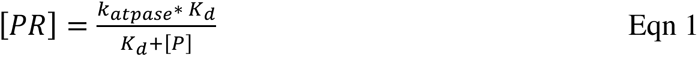

### Fluorescence intensity or anisotropy titrations to measure RNA binding

Fluorescence intensity and anisotropy measurements were performed using a FluoroMax-4 spectrofluorimeter (Horiba Jobin Yvon) in RNA Binding Buffer. NS1B constructs were titrated against a fixed amount of fluorophore-labeled RNAs (20 nM) at 25 °C. Fluorescein was excited by 495 nm light, and the emission was recorded at 515 nm. The fluorescence intensity change or fluorescence anisotropy change (F) from the initial fluorescence intensity or anisotropy (F_0_) are both proportional to the amount of protein:RNA complex formed (PR) modified by a coefficient of complex formation (f_c_), Eqn. 2. The observed change in fluorescence intensity or anisotropy was measured and plotted against protein concentration (P) and fitted to either a hyperbolic equation (Eqn. 3), a double hyperbolic equation for reactions with two phases (Eqn. 4), or a Hill equation for reactions that showed significant cooperativity (Eqn. 5) to obtain an apparent equilibrium dissociation constant (K_d_), and Hill coefficient (*n*), where applicable. Reported K_d_ values were consistently observed in titrations repeated 2 times (n = 2).

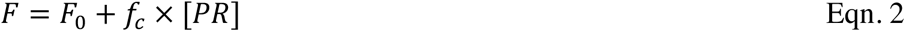

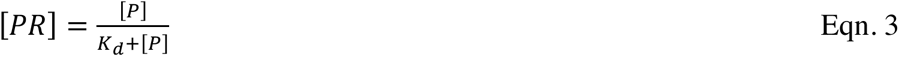

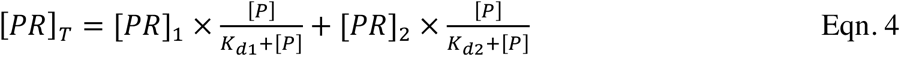

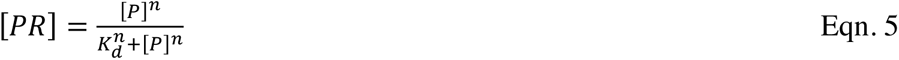

### RNA sample preparation for structural studies

HPLC-purified complementary RNA oligonucleotides [5’OH-CCAUCCUCUACAGGCG-3’ (sense) and 5’OH-CGCCUGUAGAGGAUGG-3’ (antisense)] for structural studies were purchased from Integrated DNA Technologies, Inc. (IDT). 16-bp dsRNA was prepared by mixing equal amounts of sense and antisense RNAs in an 1.5 ml Eppendorf tube which was then incubated at 94 °C for 2 minutes and gradually cooled on the bench-top at room temperature. Final buffer composition for annealing was 25 mM NH_4_OAc, pH 5.5, 275 mM NaCl, 5 mM CaCl_2_, 1 mM EDTA, 1 mM TCEP, 25 mM arginine, and 0.02% NaN_3_. RNA concentrations were determined by UV absorbance following hydrolysis of RNA. Dilute samples are incubated for 1 hour in 1 M NaOH at 37°C. The pH was then neutralized by adding equal amount of 1 M HCl. Water was used to blank and the absorbance was taken. The extinction coefficient was calculated by adding the extinction coefficients for the individual nucleic acids.

### Circular dichroism (CD) measurements

NS1B-CTD* samples were prepared for CD spectroscopy by buffer exchange into 1K buffer (40 mM NH_4_OAc, pH 5.5, 225 mM NaCl, 5 mM CaCl_2_, 0.02% NaN_3_, 50 mM arginine), addition of 1:1 ratio of RNA, and adjusting to ∼ 20 μM protein and RNA concentrations. Hence, measurements were made at significantly higher concentration than the NS1B-CTD* : 16 bp dsRNA (K_d_ ∼ 130 nM (Ma *et al*., 2016)) and NS1B-CTD* : 5’ppp HP ds10 hpRNA (K_d_ ∼ 10 nM) complex dissociation constants in this buffer. Samples were kept on ice until measurements were made. CD measurements were carried out using an AVIV Model 420 CD spectrometer and 1-mm pathlength CD cell. The scan was started at 310 nm and ended at 185 nm, taking measurements at every 2 nm. Each reading was measured for a total of 10 secs. For all samples the CD dynode was monitored to assess whether a measurement had high error. All data shown had a dynode recording between 100 mA and 600 mA. A blank of the same buffer used for buffer exchange in preparing the sample was subtracted from the sample to give final reported values, and the data are reported as molar ellipticity (mdegree / mole). Reliable CD data below 220 nm could not be obtained due to the absorbance of 50 mM L-Arg, which was required to stabilize NS1B-CTD* samples.

### Crystallization and data collection

Crystallization screening was performed using a microbatch-under-oil crystallization method at 4 °C (Luft et al., 2011) at the Hauptman Woodward Research Institute National High-Throughput Crystallization Center and in sitting drop format at Rensselaer Polytechnic Institute. Crystallization hits were the optimize in hanging drop or sitting drop experiments. dsRNA crystals useful for structure determination were obtained from mixtures of 16-bp dsRNA with NS1B-CTD* grown in drops composed of 1.0 μL of protein and 1.25 - 1.5 μL of precipitant solution [2.4 M ammonium sulfate (Hampton research HR2-541), 0.1 M bicine, pH 9.0 (Hampton research HR2-723)] under paraffin oil (Hampton Research HR3-421). A crystal grown under this condition was cryoprotected with 15% ethylene glycol prior to flash-freezing in liquid nitrogen for data collection. However, crystals similar in shape and space group also were grown from hanging drop setup (2.0 – 2.3 M ammonium sulfate, pH range from 7.5 – 9.8). All attempts to grow crystals from samples containing dsRNA alone or as a complex with NS1B-CTD* protein failed ; we observed only precipitate.

Data was collected using a Rigaku/MSC MicroMax-007 HF X-ray generator equipped with an Rigaku R-AXIS IV++ image plate detector at the Center for Advance Biotechnology and Medicine, Rutgers University. 60 frames were collected with on oscillation range 1 degree. The diffraction data from the single dsRNA crystal was processed with the HKL2000 package (Otwinowski and Minor, 1997) to 2.26 A resolution. Initial phases for 16-bp dsRNA were obtained by molecular replacement MOLREP (Vagin and Teplyakov, 2010) using U-helix 16-mer dsRNA (PDB entry 3nd3) as an initial model. The model was rebuilt using iterative cycles of manual rebuilding in Coot (Emsley and Cowtan, 2004), and was refined against 2.26 Å data with the program PHENIX (Liebschner *et al*., 2019). The data processing and refinement statistics are summarized in **Supplementary Table S1**. Atomic coordinates and structure factors were deposited in the Protein Data Bank under accession code 5KVJ.

### NMR sample preparation and data collection

Samples of NS1B-CTD* and NS1B-CTD* : 5’ppp-ds10 HP RNA complex for NMR studies were prepared in 1.7-mm or 4-mm Shigemi NMR tubes in 1K buffer (40 mM NH_4_OAc, 225 mM NaCl, 5 mM CaCl_2_, 0.02% NaN_3_, 50 mM arginine, 10% ^2^H_2_O, pH 5.5) at ∼ 100 µM protein concentration. Samples also contained trace amounts of sodium trimethylsilylpropanesulfonate (DSS) for internal chemical shift referencing. In this 1K buffer used to form a tight complex with 5’ppp-ds10 HP RNA substrate, the apo NS1B-CTD* aggregates to some degree - resulting in broader peaks in the NMR spectra for both the apo and complex samples compared to those observed in higher salt buffers (e.g. in Ma et al. (Ma *et al*., 2016)). For this reason, these samples provided less precise values of resonance frequencies, requiring a relatively large 0.065 ppm cutoff for a significant CSP (**Supplementary Figure S6**).

Backbone resonance assignments (^15^N, ^13^Cα, ^13^C’) for apo NS1B-CTD and the NS1B-CTD : 16-bp dsRNA complex, and CSPs upon complex formation, have been reported previously (Ma *et al*., 2016). NMR data for chemical shift perturbation studies of apoNS1B-CTD* vs the NS1B-CTD* : 5’ppp-ds10 HP RNA complex were collected on a Bruker Avance III 600 MHz NMR spectrometer equipped with a 5-mm triple resonance TCI cryogenic probe. NMR data were processed using NMRPipe 2.1 (Delaglio *et al*., 1995), and analyzed using SPARKY 3.106 data analysis software (Lee *et al*., 2015). Proton chemical shifts were referenced to DSS, while ^13^C and ^15^N chemical shifts were referenced indirectly using the gyromagnetic ratios of ^13^C:^1^H (0.251449530) and ^15^N:^1^H (0.101329118), respectively. Backbone resonance assignments (^15^N, ^13^Cα, ^13^C’) for apo NS1B-CTD* and NS1B-CTD* : 5’ppp ds10 HP RNA complex were determined by combined analysis of HNCO and HNCA triple resonance spectra, guided by the published assignments (Ma *et al*., 2016) for apo NS1B-CTD*.

### HADDOCK docking

Methods for using the HADDOCK server for docking protein and nucleic acid molecules have been described previously (Dominguez *et al*., 2003; van Dijk and Bonvin, 2010; van Zundert *et al*., 2016). HADDOCK takes as inputs the atomic coordinates of the protein and RNA to be docked, along with a list of the active and passive amino acids/nucleotide residues. These interactions are treated as ambiguous distance restraints to drive the docking. Active and passive residues of NS1B-CTD* were defined primarily from CSP data, as summarized in **Supplementary Table S2**. Coordinates for 16-bp dsRNA were from the X-ray crystal structure determined in this study (pdb_id 5KVJ; 2.26 Å resolution ), while the atomic coordinates for 5’ppp HP10 RNA bound to RIG-I (pdb_id 5F9H; 3.10 Å resolution) and for NS1B-CTD* (pdb_id 5DIL; 2.01 Å resolution) were published previously (Devarkar *et al*., 2016; Ma *et al*., 2016).

For docking 5’ppp ds10 HP RNA, special modifications to the RNA file were needed. HADDOCK cleans the PDB files it is given by removing any atoms it does not recognize. In the PDB file it was necessary to remove the bond between the triphosphate and the RNA. However, now that the triphosphate was no longer tethered to the RNA it is free to “float away” in the docking calculations. To overcome this, we also provided a constraint file to fix the triphosphate within 1.6 Å +/- 0.4 Å of the 5’ O5’ atom of the RNA. In CNS format: assign (resid 0 and name PG) (resid 1 and name O5’) 1.6 0.4 0.4. This provided an effective chemical bond between the triphosphate and the RNA.

In modeling the 16-bp dsRNA complex, every nucleotide of the RNA was assumed to be a possible active nucleotide because there are no data defining which specific nucleotides are involved in binding. For the 5’ppp ds10 HP RNA complex, a comprehensive sampling of possible poses was generated by defining different sections of 5’ppp ds10 HP RNA as active: 1) blunt-end active, 2) HP loop active, and 3) both blunt-end and HP loop active. In modeling each complex, 1,000 decoys were generated per simulation, clustered, and the top 4 scoring models from each cluster from all simulations were combined resulting in various decoys with a wide range of poses and interactions between NS1B-CTD* and bound dsRNA molecules. The resulting collection of coordinates for all of the resulting HADDOCK poses were combined in a single file for subsequent filtering against the SAXS data.

### Initial small angle X-ray scattering (SAXS) study of apo NS1B-CTD*

For SAXS studies of apo NS1B-CTD*, the protein was prepared in 2K buffer in a 96-well plate and SAXS was performed at the SIBYLS: beamline 12.3.1 (Advanced Light Source – Lawrence Berkeley National Laboratory, Berkeley CA). A concentration dependent SAXS study (10 mg / ml, 6 mg / ml, 4 mg / ml, 3 mg / ml, 2 mg / ml, and 1 mg / ml) determined an optimum concentration for NS1B-CTD* as 4 mg / ml (250 uM). All of the experiments at SIBYLS and other beamlines were therefore performed using this protein concentration. The SAXS data was analyzed using a combination of Scatter (Hopkins *et al*., 2017) for general SAXS curve viewing and envelope calculations, and PRIMUS (Manalastas-Cantos *et al*., 2021) for Guinier calculations, as well as P(r), R_g_, D_max_ analyses.

### SAXS study of NS1B-CTD bound to 16-bp dsRNA

Initial SAXS experiments were used to determine optimal concentrations, conditions, and NS1B-CTD* : 16-bp dsRNA ratios for best signal-to-noise, and to also minimize aggregation, and a sample concentration dependence study was performed prior to production data collection. Optimal conditions for SAXS experiments were identified, and samples of NS1B-CTD* were prepared at 250 μΜ (4 mg/mL), after 16-bp dsRNA was added to a 1:1 NS1B-CTD : 16-bp dsRNA ratio in 2K buffer (**Supplementary Table S4**). To minimize aggregation in the SAXS samples, samples were centrifuged at 8,500 x g for a minimum of 10 mins immediately prior to measurements. For this complex, SAXS data was collected on CHESS beamline G1 (MacCHESS: 161 Synchrotron drive, Ithaca NY – Cornell University) at 9.924 keV (1.249 Å) with 7.7×1011 photons/s. The X-ray beam was collimated to 250 × 250 µm^2^ diameter and centered on a capillary sample cell with 1.5 mm path length and 25 μm thick quartz glass walls (Charles Supper Company, Natik, MA). The sample cell and full X-ray flight path, including beamstop, were kept *in vacuo* (< 1×10-3 Torr) to eliminate air scatter. Temperature was maintained at 4 °C. Images were collected on a dual Pilatus 100K-S detector system (Dectris, Baden, Switzerland). Sample-to-detector distance was calibrated using silver behenate powder (The Gem Dugout, State College, PA). The useful q-space range (4πSinθ/λ with 2θ being the scattering angle) was generally from q_min_= 0.01 Å^-1^ to q_max_= 0.27 Å^-1^. Image integration, normalization, and subtraction was carried out using the BioXTAS RAW program (Hopkins *et al*., 2017). Radiation damage was assessed using the CorMap criterion (Hopkins *et al*., 2017)as implemented in RAW’s built-in averaging function. Sample and buffer solutions were normalized to equivalent exposure before subtraction, using beamstop photodiode counts. Sample plugs of approximately 20-30 μl were delivered from a 96-well plate to the capillary using a Hudson SOLO single-channel pipetting robot (Hudson Robotics Inc. Springfield, New Jersey). To reduce radiation damage, sample plugs were oscillated in the X-ray beam using a computer-controlled syringe pump. Typically, 10-20 undamaged 1s exposures were averaged to produce buffer and sample profiles. Scattering intensities were placed on an absolute scale using water as a standard. SAXS data quality was assessed by Guinier plot analysis (**Supplementary Figure S2A**). These data indicated little or no aggregation, low sample radiation damage, and appropriate P(r) curve shapes.

### SAXS study of NS1B-CTD* bound to 5’ppp ds10 HP RNA

Samples of NS1B-CTD* were prepared at a concentration of 250 μΜ (4 mg/mL) and 1 : 1 ratio of NS1B-CTD* : 5’ppp ds10 HP RNA in 1K buffer (**Supplementary Table S4**), since this lower salt buffer provides a tighter complex. Immediately before mixing the protein with RNA to make the complex, NS1B-CTD* was buffer exchanged into 1K buffer, and concentrated to provide a 250 μΜ complex concentration after adding RNA. RNA was then added to provide a 1:1 NS1B-CTD : 5’ppp ds10 HP RNA ratio. Samples were centrifuged at 8,500 x g for a minimum of 10 mins at the beamline immediately before data collection on the supernatant. SAXS was performed at BioCAT beamline 18ID (Advanced Photon Source, Chicago, IL). Data were collected in a SAXS flow cell which consists of a 1.5 mm ID quartz capillary with 10 µm walls held at 20 °C using 12 keV incident X-rays. Scattering intensity was recorded using a Pilatus3 × 1M (Dectris) detector which was placed 3.44 m from the sample, providing access to a q-range of 0.012 Å^-1^ to 0.3 Å^-1^. During exposure, sample was flowed through the beam in a single direction, and 0.5 s exposures were acquired every 2 seconds during flow. Data was reduced using BioXTAS RAW 1.4.1. Buffer blanks were created by recording SAXS data from a matched buffer and subtracted from exposures from the sample to create the I(q) vs q curves used for subsequent analyses. All sets of nominally identical measured profiles were automatically compared using CorMap (Hopkins *et al*., 2017) to test for radiation damage or other changes. SAXS data quality assessed by Guinier plot analysis (**Supplementary Figure S2B**). These data indicated little or no aggregation of the complex under these conditions, consistency between expected and observed molecular weight, low sample radiation damage, and appropriate P(r) curve shapes.

### SAXS data analysis

A combination of RAW (Hopkins *et al*., 2017), Scatter (Hopkins *et al*., 2017), and PRIMUS (Manalastas-Cantos *et al*., 2021) software packages were used for SAXS data analysis. In addition, comparison of experimental SAXS data and those predicted from models was done using FOXS webserver (Schneidman-Duhovny *et al*., 2016). Calculated SAXS curves for models were then downloaded, and P(r), R_g_, and D_max_ were calculated using PRIMUS (Manalastas-Cantos *et al*., 2021). To compare the P(r) plots it was required to export the data points from the GNOM file that is created when calculating the P(r), into an Excel spreadsheet, and create a plot within Excel. Visualizations of models and SAXS envelopes were done using UCSF ChimeraX modeling software (Pettersen *et al*., 2021). The combined NMR, RNA X-ray crystal structures, and CD data were used to guide integrative modeling of the NS1B-CTD : 16-bp dsRNA and NS1B-CTD : 5’ppp-ds10 HP RNA complexes using HADDOCK (Dominguez *et al*., 2003; van Dijk and Bonvin, 2010; van Zundert *et al*., 2016). All graphical analysis and structural model figures were created using UCSF ChimeraX (Pettersen *et al*., 2021).

### Viruses and cells

B/Yamanashi/166/98 NS1 mutant viruses were generated by 8-plasmid reverse genetics (Hoffmann et al., 2002), and all 8 gene segments of the viruses were sequenced to verify that only the correct mutation was generated. Mutant and wild-type viruses were grown in 9-day-old fertilized chicken eggs. The titers of virus stocks were determined by plaque assays in MDCK cells. For viral infections, A549 cells in 60 mm dishes were infected with 2.0 plaque-forming units (p.f.u.) per cell of the indicated virus. After incubation at 34 °C for one hour, the viral inoculum was removed, and the infected cells were further incubated in OPTI-MEM medium for 2, 4, 6, or 8 hours. Infected cells were collected, freeze-thawed three times (Robitaille et al., 2016), and then extracted using the PhosphoSafe reagent (Novagen) (Kuo *et al*., 2010). An aliquot (20 μg) of each extract was subjected to SDS-gel electrophoresis, followed by immunoblotting with the indicated antibodies. The rabbit monoclonal antibodies against IRF-3, phosphor-IRF-3 (Ser396) (#4947) and RIG-I (#3743) were purchased from Cell Signaling Technology. The mouse monoclonal antibody against NP of influenza B virus was purchased from Southern Biotech, and the rabbit antibody against the NS1B protein was generated as previously described (Zhao et al., 2016).

## QUANTIFICATION AND STATISTICAL ANALYSIS

Quantification of ATP hydrolysis assays was performed with ImageQuant TL 8.1 software (Cytiva Life Sciences). Fitting ATP hydrolysis assays, fluorescence intensity assays, and fluorescence anisotropy assays was done using SigmaPlot v11.0. Associated errors of measurements of fitting and number of sample sets is noted in corresponding figure legends.

## SUPPLEMENTARY TABLES

**Table S1.** 16-bp RNA X-ray crystallography diffraction data and refinement statistics^a^.

|  |  |
| --- | --- |
| **Crystal Parameters** |  |
| Space group | H 3 |
| Cell dimensions |  |
| a, b, c (Å) | 43.3, 43.3, 124.4 |
| $\alpha$, $\beta$, $\gamma$, (°) | 90, 90, 120 |
| Matthews coefficient (Å<sup>3</sup>/Da) | 2.13 |
| Solvent Content (%) | 42.15 |
| **Data Collection<sup>b</sup>** |  |
| Wavelength (Å) | 1.542 |
| Resolution (Å) | 21.7-2.26 (2.30-2.26) |
| R<sub>sym</sub> (%) | 2.8 (33.4) |
| No. of unique reflections | 3824 |
| No. of reflections in R<sub>free</sub> set | 189 |
| Mean redundancy | 2.0 (1.7) |
| Overall completeness (%) | 93.6 (97.7) |
| Mean I/$\sigma$ | 37.2 (3.5) |
| **Refinement Residuals<sup>c</sup>** |  |
| R<sub>free</sub> (%) | 23.3 (34.0) |
| R<sub>work</sub> (%) | 19.4 (34.3) |
| Completeness (%) | 93.1 (96.4) |
| **Model Quality** |  |
| RMSD<sup>a</sup> bond lengths (Å) | 0.010 |
| RMSD<sup>a</sup> bond angles (°) | 1.34 |
| dsRNA B-factor (Å<sup>2</sup>) | 45.0 |
| Mean overall B-factor<br>(dsRNA + solvent) (Å<sup>2</sup>) | 45.0 |
| Mean solvent B-factor (Å<sup>2</sup>) | 54.9 |
| **Model Contents** |  |
| Protomers in ASU<sup>a</sup> | 1 |
| Nucleic Acids | A1-16; B1-16 |
| Ligand | Arg |
| No. of dsRNA atoms | 682 |
| No. of heterogen atoms | 0 |
| No. of water molecules | 10 |

<sup>a</sup> Standard definitions were used for all parameters (Drenth, 1984). Entries in parentheses report data from the limiting resolution shell. Data collection and refinement statistics come from *SCALEPACK* (Otwinowski and Minor, 1997) and *PHENIX* (Liebschner *et al.*, 2019)], respectively. The abbreviations RMSD and

**Table S2.** Residues treated as Active or Passive in HADDOCK based on Chemical Shift Perturbation (CSP) NMR data. Residues defined as “Active or “Passive” for HADDOCK modeling of complexes formed between NS1B-CTD* and either 16-bp dsRNA or 5’ppp ds10 HP RNA. Active residues for 16-bp dsRNA are those with CSPs greater than 30 ppb while passive residues are those with CSPs between 30 ppb and 20 ppb. Active residues for 5’ppp ds10 HP are those with CSPs greater than 95 ppb and passive residues are those with CSPs between 95 ppb and 65 ppb. For the 5’ppp-ds10 HP RNA complex, CSPs for residue Lys 221 could not be measured, but it was treated as “active” because mutation to Ala affects its RNA binding affinity. Active nucleic acid residues for docking were defined as explained in the Methods section.

| HADDOCK residue type | active NS1B-CTD residues for 16-bp dsRNA complex | active NS1B-CTD residues for 5’ppp-ds10 HP |
| --- | --- | --- |
| Active binding | 144, 145, 147, 148, 150, 152, 153, 155, 158, 159, 161, 162, 164, 170, 189, 192, 193, 195, 202, 204, 205, 206, 209, 221, 222, 223, 225, 226, 227, 228 | 143, 144, 145, 147, 150, 152, 157, 159, 160, 169, 171, 178, 179, 193, 195, 196, 204, 208, 209, 237, 241, 259, 260, 264, 265 |
| Passive binding | 146, 149, 156, 157, 160, 171, 172, 194, 211, 242, 252, 257, 262, 264, 266, | 151, 158, 161, 163, 166, 167, 170, 173, 176, 188, 190, 191, 202, 203, 212, 224, 226, 227, 228, 229, 258, 263, 266, 268 |
| No CSP data | 169, 178, 182, 185, 190, 208, 218, 224, 232, 234, 237, 248, 255, 259, 263, 267, 268, 272, 273, 274, 275, 281 | 153, 155, 156, 162, 174, 197, 205, 206, 207, 221, 225, 254, 261, 276 |

**Table S3.** Small angle X-ray scattering (SAXS) data.

| SAXS Data Type | 16-bp ds RNA :<br>NS1B-CTD* Complex | 5'ppp ds10 HP:<br>NS1B-CTD* Complex |
| --- | --- | --- |
| Beamline | Cornell – CHESS | BIOCAT |
| Porod-Debye (Px) | 2.7 | 3.1 |
| Low q ($\text{\AA}^{-1}$) | 0.0063 | 0.012 |
| High q ($\text{\AA}^{-1}$) | 0.275 | 0.299 |
| Reciprocal $R_g$ ($\text{\AA}$) | 28.11 | 20.48 |
| Real Space $R_g$ ($\text{\AA}$) | 28.21 | 20.35 |
| $D_{\text{max}}$ ($\text{\AA}$) | 103 | 69 |
| Energy (keV) | 9.924 | 12.00 |
| Temperature ($^{\circ}\text{C}$) | 20 | 20 |
| Exposure | 1 sec x 10 sec | 0.5 sec / 2.0 sec |
| Concentration of complex<br>(mg / mL) | 4.00 | 4.00 |

**Table S4.** Aqueous buffers used in sample preparation and biophysical studies.

| Buffer | Components and concentrations |
| --- | --- |
| BB | 50 mM Tris, pH 7.5, 500 mM NaCl, 1 mM TCEP, 40 mM imidazole, 0.02% $\text{NaN}_3$. |
| EB | 50 mM Tris, pH 7.5, 500 mM NaCl, 1 mM TCEP, 500 mM imidazole, 0.02% $\text{NaN}_3$. |
| LS | 10 mM Tris, pH 7.5, 100 mM NaCl, 0.02% $\text{NaN}_3$. |
| A | 50 mM Tris, pH 8.0, 5 mM DTT. |
| B | 50 mM Tris, pH 8.0, 1 M NaCl, 5 mM DTT. |
| 2K | 40 mM $\text{NH}_4\text{OAc}$, pH 5.5, 450 mM NaCl, 5 mM $\text{CaCl}_2$, 0.02% $\text{NaN}_3$, 50 mM arginine. |
| 1K | 40 mM $\text{NH}_4\text{OAc}$, pH 5.5, 225 mM NaCl, 5 mM $\text{CaCl}_2$, 0.02% $\text{NaN}_3$, 50 mM arginine. |
| FL | 50 mM Tris, pH 8.0, 300 mM NaCl |
| RNA Binding<br>Buffer | 50 mM MOPS, pH 7.4, 5 mM DTT, 5 mM $\text{MgCl}_2$, 0.01% Tween20. |
| Storage Buffer | 20 mM Na-phosphate, pH 7.7, 1 mM $\text{Na}_3\text{-EDTA}$, 1 mM DTT, 100 mM NaCl, 50% (v/v) glycerol. |

## SUPPLEMENTARY FIGURES

**Fig. S1.**
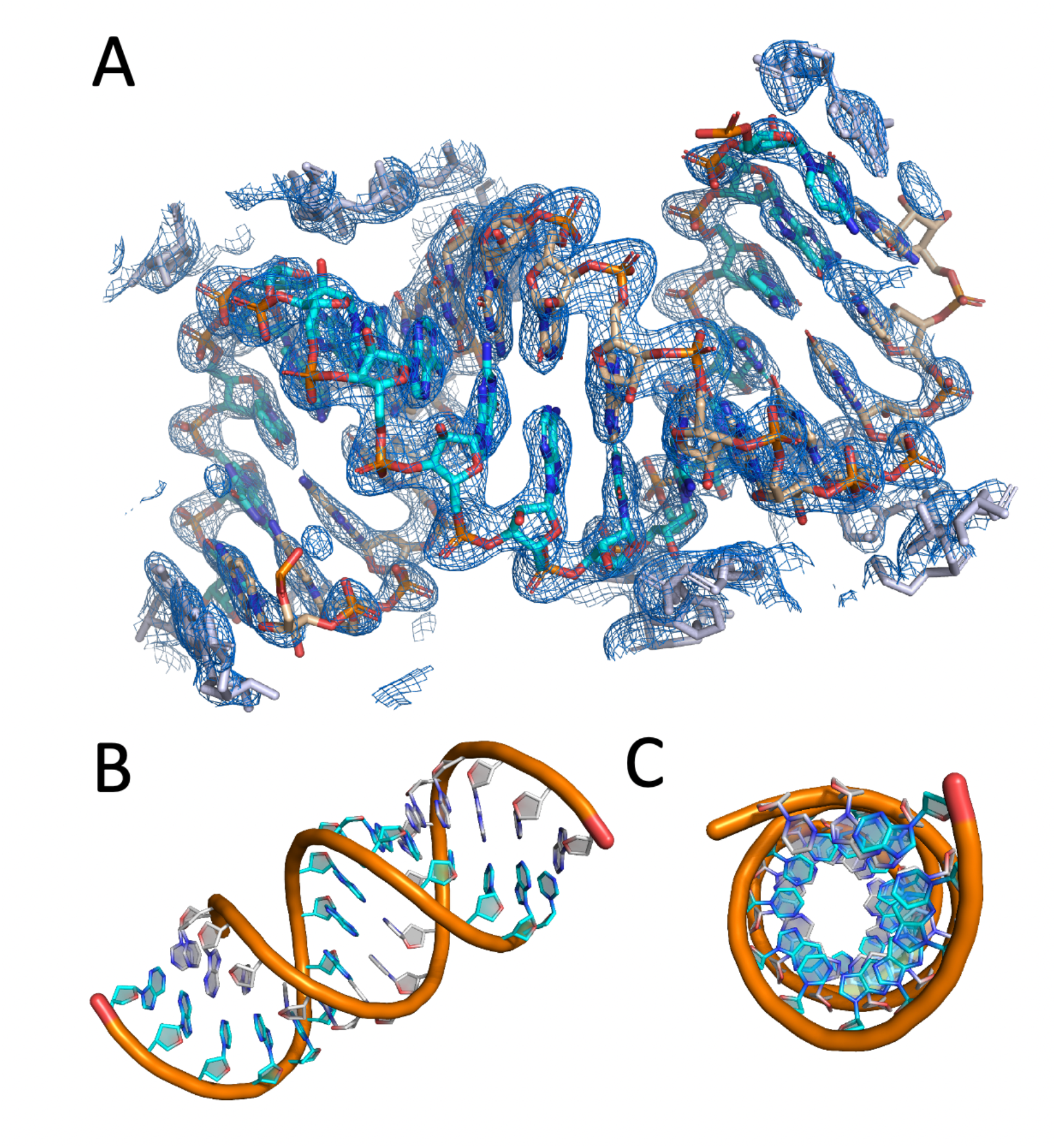
X-ray crystal structure of 16-bp dsRNA. (A) Rendering of the 2.26 Å resolution 2F_o_-F_c_ electron density map. Electron density map has been contoured to 1.5 σ. Carbons from RNA chain A and B have been rendered in teal and beige respectively. Symmetry related models have been rendered in gray. (B) Side-view cartoon representation of the double stranded A-form RNA helix. (C) end-on view of the RNA helix. Coordinates and structure factor data are deposited in the Protein Data Bank as entry 5KVJ. Related to Figure 1

**Fig. S2.**
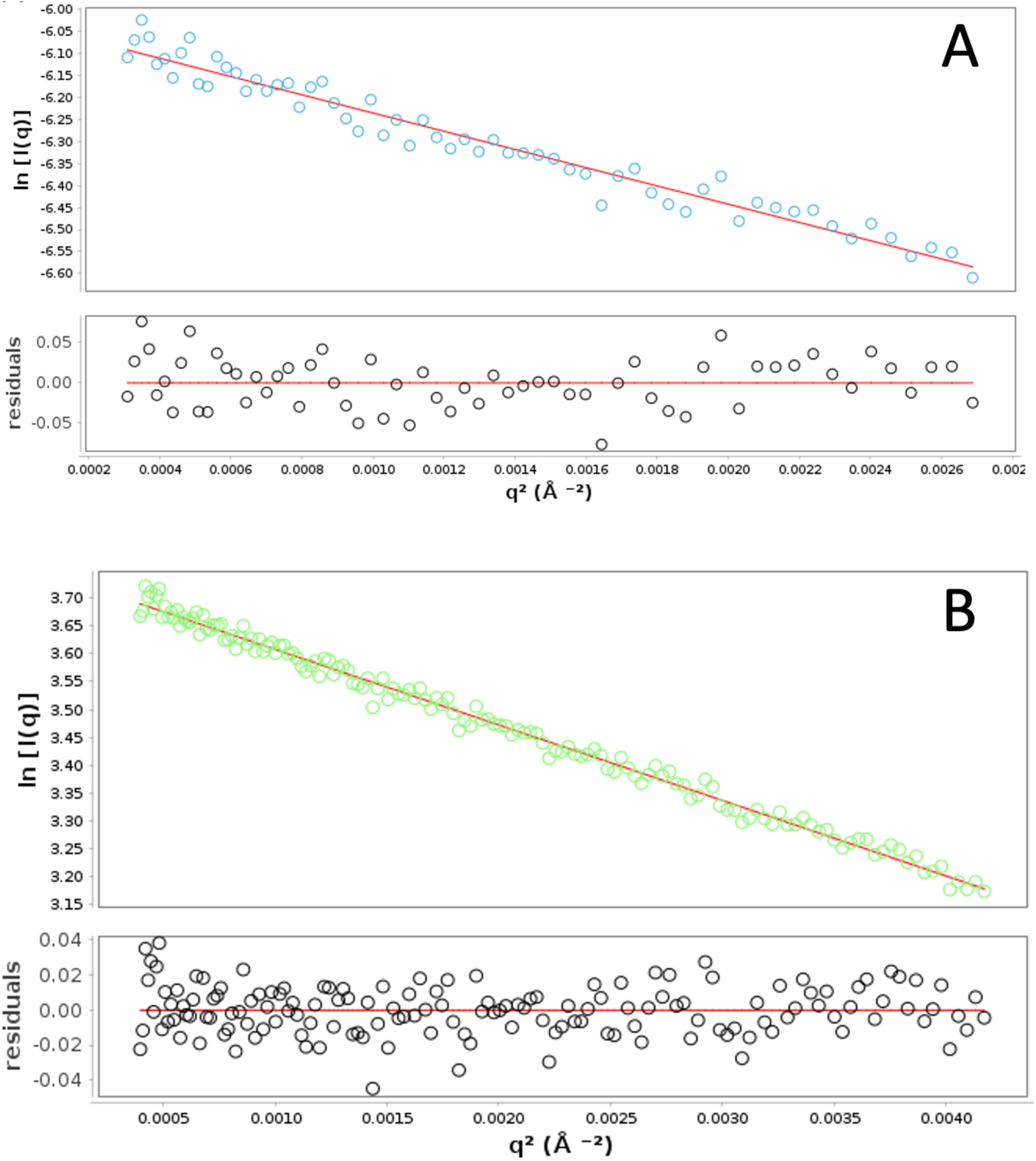
Guinier plots for NS1B-CTD : RNA complexes. (A) dsRNA : NS1B-CTD* at a ratio of 1 : 1, concentration of 250 μΜ, and 4 °C. (B) 5’ppp ds10 HP : NS1B-CTD* at a ratio of 1 : 1, concentration of 250 μΜ, and 20 °C. Linear regions were plotted by trimming data to no less than 50 data points, up to the point where the data became non-linear. In each panel, the top graph shows Guinier fitting plot and the bottom graph shows linear residuals. Related to Figures 1 and 3.

**Fig S3.**
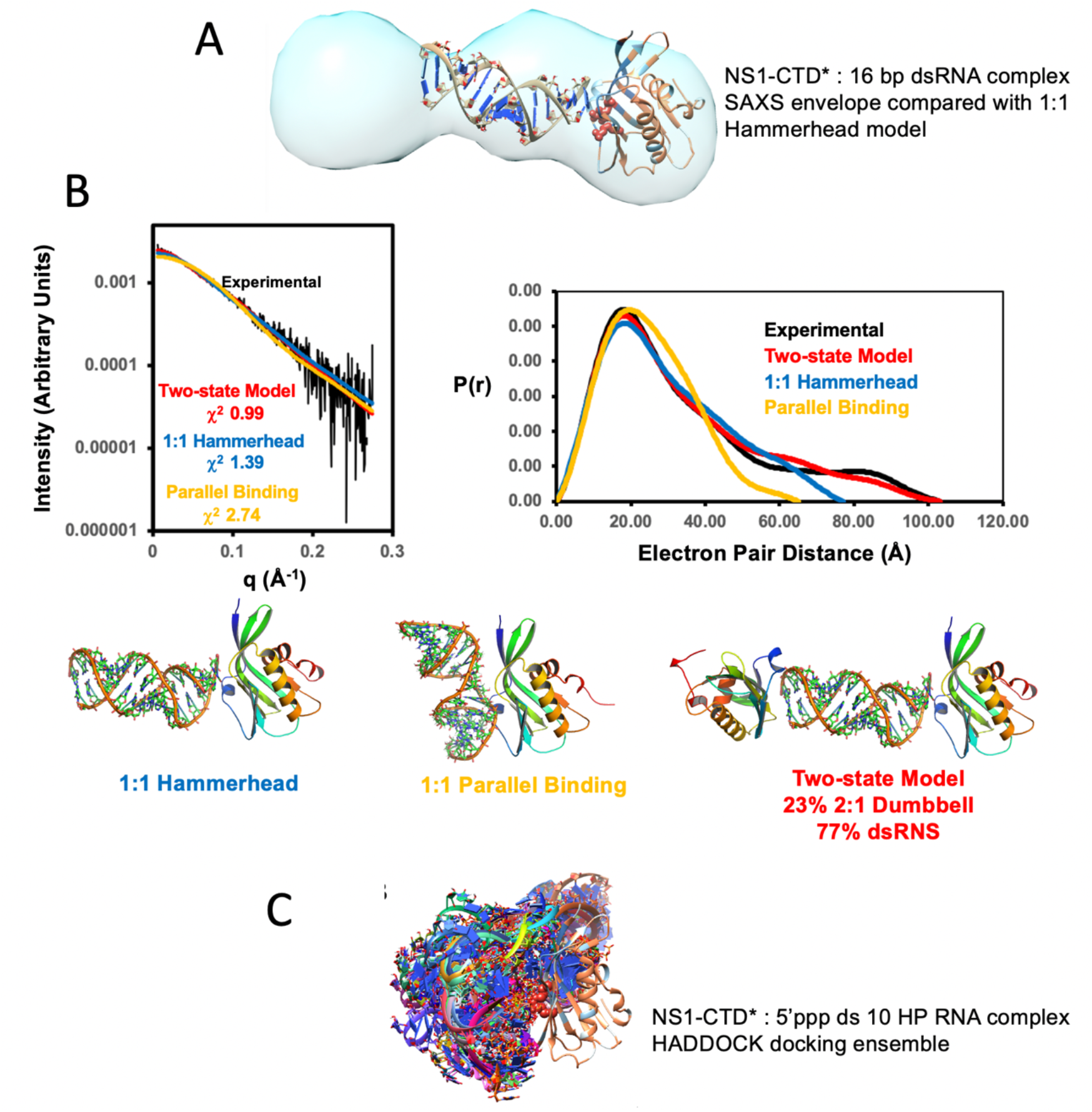
Supporting details of SAX studies. (A) SAXS envelope consistent with NS1B-CTD end binding to 16-bp dsRNA. (A) SAXS envelope for NS1B-CTD* : 16-bp dsRNA complex revealed an extended structure with blunt-end binding, as opposed to one where the protein is bound to the middle of the RNA helix in a parallel orientation as observed for the NS1B-NTD : dsRNA complex (Yin *et al*., 2007). The SAXS envelope is shown together with the best-fit model from the HADDOCK docking result. The density also suggests the complex may have NS1B-CTD* bound at both ends of the dsRNA helix. **(B) SAXS data exclude parallel binding mode and indicate a Two-state Model of a mixture of a ternary complex and free dsRNA**. Comparison of SAXS data fits for two-state model (red), 1:1 hammerhead (blue), and best fitting parallel-binding model (yellow) for NS1B-CTD* : dsRNA complex. The reciprocal (left) and real (right) space scattering data are compared with data back-calculated from the three models shown at the bottom of this panel. **(C) Models of NS1B-CTD* : 5’ppp ds10 HP RNA complex generated by HADDOCK prior to SAXS filtering**. The models are oriented with the NS1B-CTD overlaid to show the variation in sampling of relative orientations relative to the RNA. This ensemble was then filtered against the SAXS data using the FoXS server. Coloring for each model matches coloring scheme of Figures 1 and 3. Related to Figures 1 and 3.

**Fig. S4.**
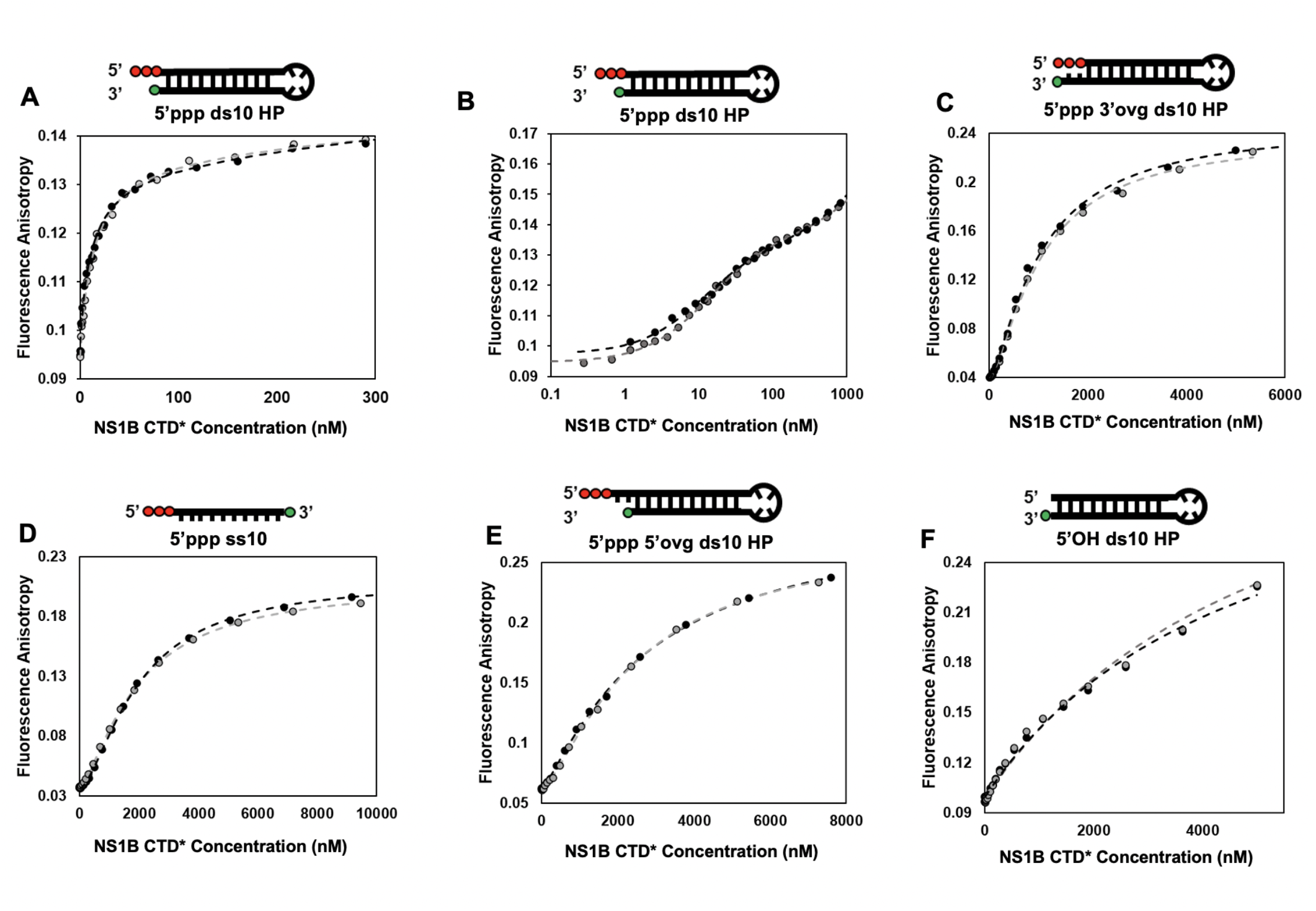
Individual anisotropy experiments for each tested RNA with NS1B CTD*. Each experiment was repeated two times on different days (n = 2). Dotted lines represent the fit for each trial. (A) 5’ppp ds10 HP RNA trials, fit to a double hyperbola. Only the first phase is shown. (B) 5’ppp ds10 HP RNA trials, fit to a double hyperbola and plotted on a logarithmic x-axis. The full profile is shown. (C) 5’ppp 3’ovg ds10 HP RNA trials, fit to a Hill equation (n=1.4 and 1.3). (D) 5’ppp ss10 trials, fit to a Hill equation (n=1.4 and 1.5). (E) 5’ppp 5’ovg ds10 HP RNA trials, fit to a Hill equation (n=1.2 and 1.4). (F) 5’OH ds10 HP RNA trials, fit to a single hyperbola. Related to Figure 2.

**Fig. S5.**
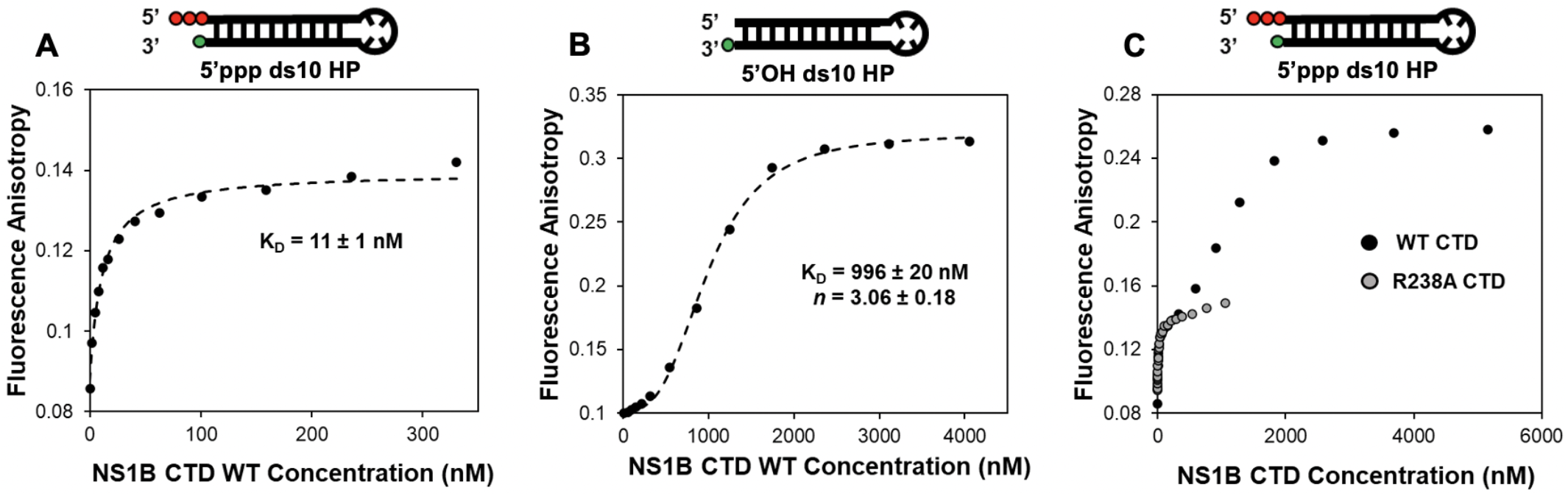
Both WT NS1B-CTD and R238A mutant NS1B-CTD* have a preference for 5’ppp ends over 5’OH ends. (A) Highlight of the first phase of WT NS1B-CTD binding to 5’ppp ds10 HP. K_d_ was determined by hyperbolic regression fitting. (B) 5’OH ds10 HP RNA binding to WT NS1B-CTD. K_d_ and Hill coefficient *n* were determined by fitting to a Hill regression plot, as the data showed significant sigmoidicity. (C) Overlay of absolute anisotropy values of both WT NS1B-CTD and R238A NS1B-CTD* binding to 5’ppp ds10 HP RNA. Each binding experiment was repeated twice (n = 2). A representative binding curve is shown. WT - wild type. Related to Figure 2.

**Fig. S6.**
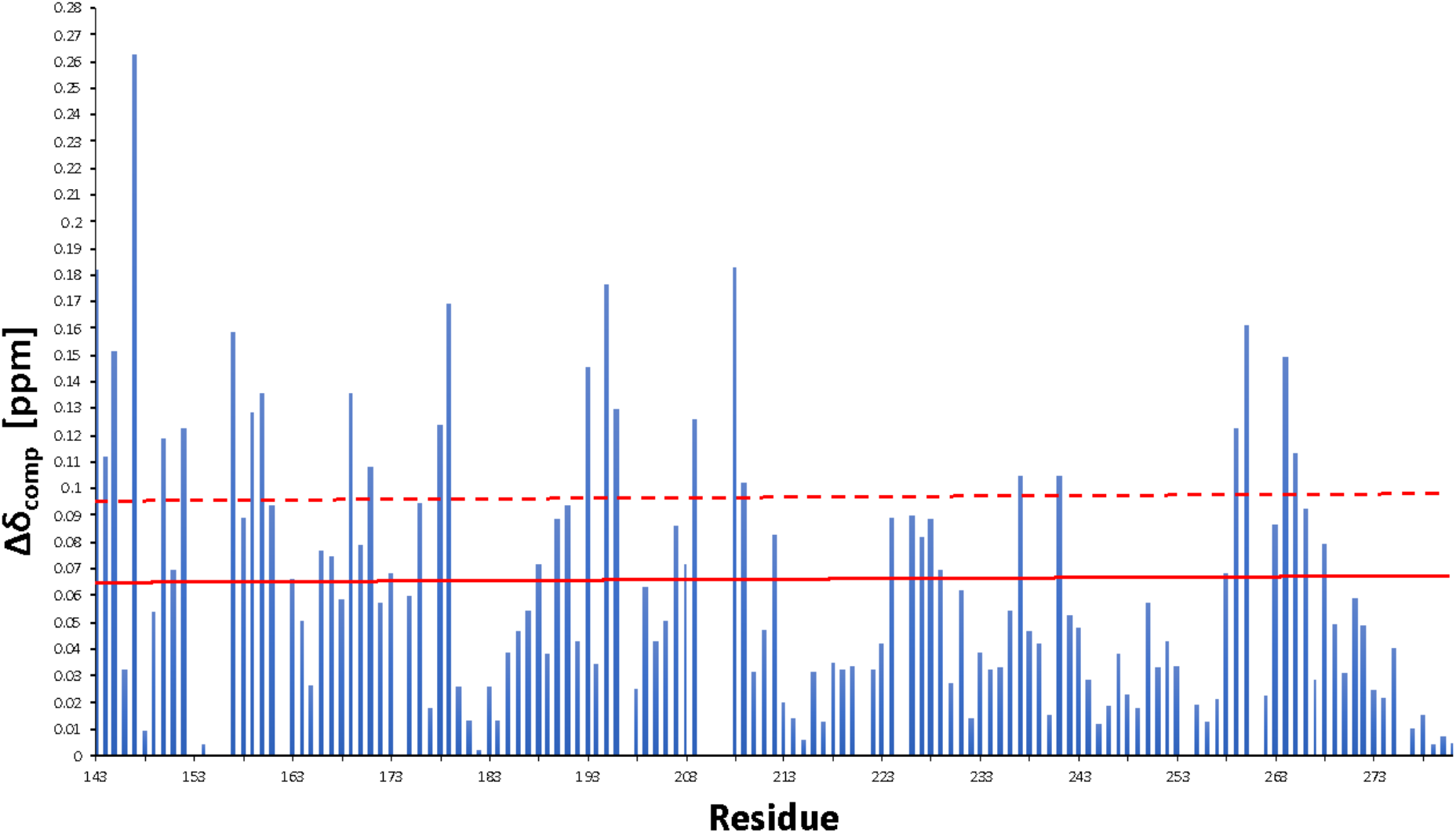
Chemical shift perturbations (CSPs) due to formation of NS1B-CTD* : 5’ppp ds10 HP RNA complex. CSPs of NS1B-CTD* due to 5’ppp ds10 HP binding are mapped along the protein sequence. The solid horizontal line at 65 ppb, and the dashed horizontal line at 95 ppb represent the cutoffs used to define passive (65 < Δδ_comp_ < 95 ppb) and active (Δδ_comp_ > 95 ppb) residues involved in RNA-binding. Related to Figure 3.

**Fig. S7.**
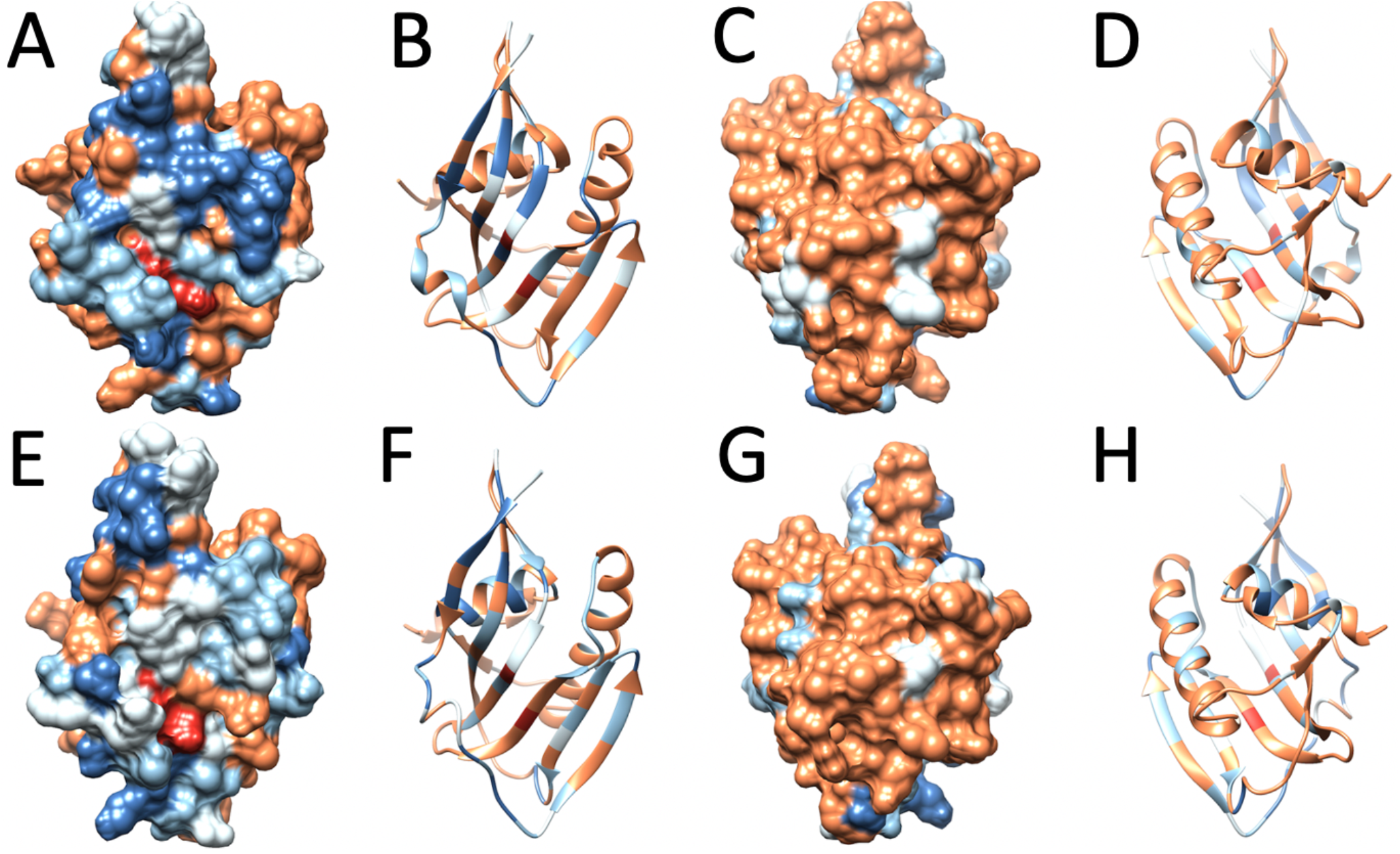
Details of chemical shift perturbation studies. **(A-D) ^15^N-^1^H CSPs due to 16-bp dsRNA binding mapped onto the 3D structure of NSB1-CTD (Ma *et al*., 2016).** Panels C and D are orientations of the molecules rotated by 180°. (**E-H) ^5^N-^1^H CSPs due to 5’ppp ds10 RNA binding mapped onto the 3D structure of NSB1-CTD**. Panels G and H are orientations of the molecules rotated by 180°. In all panels, dark blue - high CSP, light blue - medium CSP, white - no data, brown - little or no CSP. Residues Arg208 and Lys221 are colored red. Panels A,C,E,G are shown in surface representation. while panels B,D,F,H are shown in ribbon representation to better see the locations of CSPs. Related to Figures 1 and 3.

